# Long-term, super-resolution HIDE imaging of the inner mitochondrial membrane in live cells with a cell-permeant lipid probe

**DOI:** 10.1101/2022.10.19.512772

**Authors:** Shuai Zheng, Neville Dadina, Deepto Mozumdar, Lauren Lesiak, Kayli Martinez, Evan W. Miller, Alanna Schepartz

**Affiliations:** Department of Chemistry, University of California, Berkeley, CA 94720, USA; Department of Chemistry, Yale University, New Haven, CT 06511, USA; Department of Molecular and Cell Biology, University of California, Berkeley, CA 94720, USA; California Institute for Quantitative Biosciences, University of California, Berkeley, CA 94720, USA; Chan Zuckerberg Biohub, San Francisco, CA 94158, USA

## Abstract

The densely packed inner mitochondrial membrane (IMM) is vital for bioenergy generation and its dynamics control mitochondrial health and cellular homeostasis. IMM structure is complex, however, and imaging its dynamics with high temporal and spatial resolution is complicated by the photosensitivity of IMM-resident enzymes. Here we describe the cell-permeant, lipid-like acridine orange derivative MAO-N_3_ and use it to assemble high-density, environmentally sensitive (HIDE) probes that selectively label and image the IMM in live cells. MAO-N_3_ pairs with multiple SPAAC-reactive fluorophores to support HIDE imaging *via* confocal, Structured Illumination, Single Molecule Localization, and Stimulated Emission Depletion microscopy, all with significantly improved resistance against photobleaching. The HIDE probes generated using MAO-N_3_ require no genetic manipulations, are non-toxic in model cell lines and primary cardiomyocytes, even under conditions that amplify the effects of mitochondrial toxins, and visualize the IMM for up to 12.5 hours with unprecedented spatial and temporal resolution.

## Main

The mitochondrion is the powerhouse of the cell and contributes to numerous physiological processes including programmed cell death, innate immunity, autophagy, redox signaling, calcium homeostasis, and stem cell reprogramming. It powers the cell using oxidative phosphorylation, sustains the cell via multiple cell signaling activities, and kills the cell when necessary by initiating programmed cell death^1^. Mitochondria are distinguished by two distinct membranes. The outer mitochondrial membrane (OMM) acts as a diffusion barrier, transduces signals in and out of the organelle, and mediates multiple inter-organelle interactions^2^. The inner mitochondrial membrane (IMM) consists of two discrete subcompartments, the inner boundary membrane (IBM) and the cristae. The IBM parallels the OMM and contains transporters that shuttle ions and metabolites between the mitochondrial matrix (the space parameterized by the IMM, which houses enzymes responsible for the tricarboxylic acid cycle and heme biosynthesis) and the intermembrane space. The cristae are densely packed membrane invaginations with diffraction-limited dimensions that support oxidative phosphorylation, mitochondrial DNA maintenance, and iron–sulfur cluster biogenesis^1^. The IMM and OMM undergo continual fission and fusion, and individual crista remodel continuously in response to signals from other organelles^2^. These dynamic events drive mitochondrial biogenesis and recycling and play a pivotal role in apoptosis signaling^2^. Multiple pathological conditions, in both neurons (Charcot–Marie–Tooth disease)^2^ and energy-demanding myocytes (Barth’s syndrome for cardiomyocytes^3,4^ and Kearns-Sayre syndrome^5^ for eye muscles) are associated with mutations in IMM-localized enzymes. The on-demand initiation of mitochondrial apoptosis is also desirable in the development of novel cancer therapies^6^.

Imaging the IMM is challenging due to its complex and diffraction-limited dimensions, especially if the goal is to study dynamics. The respiratory chain enzymes that localize within cristae are sensitive to fluorophore-mediated or high-energy light-induced damage^7,8^ that complicates imaging using Stimulated Emission Depletion (STED)^9^ and Single Molecular Localization Microscopy (SMLM)^10,11^. Some progress in visualizing IMM dynamics at super-resolution has been achieved with small molecule dyes. MitoPB Yellow, for example, is relatively photostable, localizes selectively to the IMM, and can be combined with STED to image IMM dynamics over extended time (300 frames over 390 s)^12^. However, MitoPB Yellow has limited spectral tuning potential for multi-color imaging and longer time-lapse imaging is precluded by the requisite high-energy (488 nm) excitation wavelength–mitochondrial swelling is induced after acquisition of 150 frames. Moreover, since MitoPB Yellow is not modular, it is not easily adapted to other imaging modalities. MitoEsq-635^8^ and PK Mito Orange (PKMO)^13^ also localize selectively to the IMM and are sufficiently photostable for STED imaging. MitoEsq-635 is excited by lower energy light (633 nm) than MitoPB Yellow, but is also less photostable, supporting the acquisition of only 40 STED frames over 50 min to visualize IMM dynamics^8^. PKMO is also excited by lower energy light (591 nm) and supports two-color STED imaging, but can support only 33 frames over 5 min^13^. The study of cristae remodeling and inter-organelle crosstalk is currently precluded by the absence of probes capable of long time-lapse imaging with high spatiotemporal-resolution. The ideal probe for the IMM should be spectrally tunable, bright, photostable, and capable of supporting multiple imaging modalities.

High-density, environmentally sensitive (HIDE) probes are small molecules that selectively label organelle membranes and resist photobleaching, leading to exceptionally long time-lapse images of organelles in live cells, even at super-resolution^14–21^. Each HIDE probe consists of two parts (**Fig. 1a**). The first is an organelle-specific, lipid or lipid-like small molecule that is click-ready by virtue of a reactive functional group such as an azide (N_3_). The second is a far-red fluorophore equipped with the appropriate reaction partner. These two parts, when added sequentially to live cells, undergo a bioorthogonal reaction that localizes the fluorophore at high density within the organelle membrane (**Fig. 1b**)^17^. The combination of dense localization and a hydrophobic environment results in a 10-50-fold enhancement in apparent photostability. HIDE probes have been used for long time-lapse imaging of the ER, Golgi, plasma membrane, mitochondrial matrix, and late endolysosomes^14–21^. They label organelles in model cell lines, primary cells from patient samples^19^, even neurons^18,20^. And because they are assembled from two parts, the emission properties can be altered at will to support multi-color labeling and all established modalities^20^. One mitochondrial HIDE probe (RhoB-TCO) has been reported^17,20^, but it labels mitochondrial subcompartments with minimal specificity; it is not selective for the IMM and cannot resolve IMM dynamics and fine structure^17,20^.

**Fig. 1.**
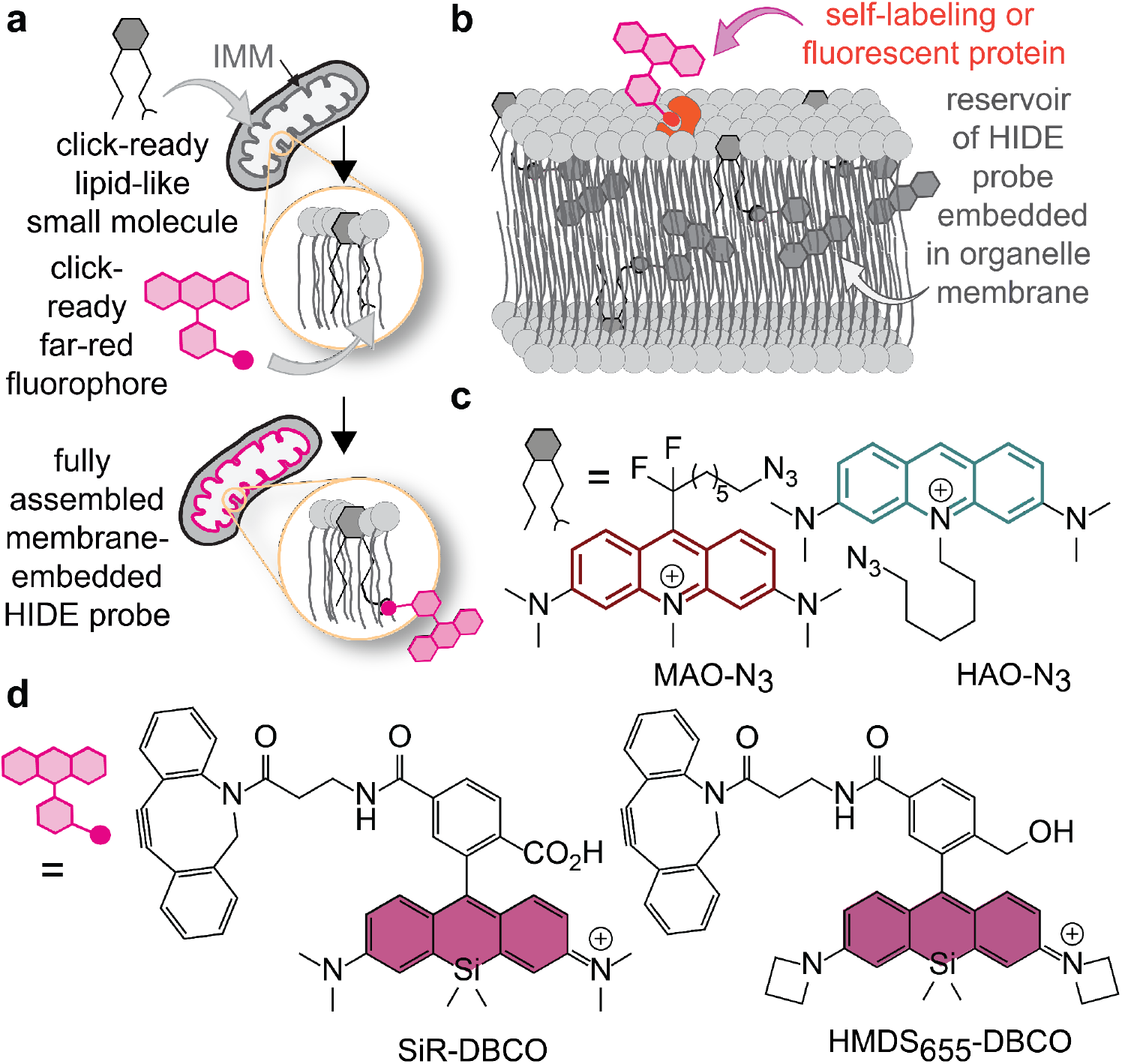
New HIDE probes to image the inner mitochondrial membrane. **a,** HIDE probes assemble from two parts. One is a lipid-like small molecule appended to a reactive bioorthogonal functional group, such as an azide (N_3_). The second is a modality-specific fluorophore equipped with the appropriate click reaction partner. **b,** HIDE probes prolong imaging time because they localize within organelle membranes at higher density than possible with a self-labeling or fluorescent protein, and because in this hydrophobic environment only a small fraction of the molecules absorb light at the excitation wavelength. Since fewer molecules absorb light at any given time, fewer molecules bleach; the result is a reservoir of fresh fluorophores that can replenish photobleached dye molecules during an imaging experiment. **c,** Derivatives of acridine orange, namely MAO-N_3_ and HAO-N_3_, used to generate the HIDE probes reported here. **d,** Structures of two silicon rhodamine dyes used in conjunction with MAO-N_3_ and HAO-N_3_. HMDS_655_-DBCO was used for single molecule localization microscopy (SMLM), whereas SiR-DBCO was used for all other imaging modalities.

Here we describe two new lipid-like small molecules, MAO-N_3_ and HAO-N_3_ (**Fig. 1c**) and use them to assemble HIDE probes selective for the inner mitochondrial membrane. These HIDE probes are largely benign to mitochondrial health, even under conditions that amplify mitochondrial toxicity. When paired with the appropriate fluorophore, MAO-N_3_ supports long time-lapse imaging of the IMM using confocal, Structured Illumination (SIM), Single-Molecule Localization (SMLM), and Stimulated Emission Depletion (STED) microscopy. When paired with SiR-DBCO (**Fig. 1d**) and visualized using confocal microscopy, MAO-N_3_ could visualize the mitochondria for more than 12.5 h with minimal loss in signal intensity or cell viability, whereas the commercial dye MitoTracker DeepRed lost > 50% signal intensity within 2h and the labeled cells proliferate slowly. When paired with SiR-DBCO and visualized using SIM, MAO-N_3_ could visualize the IMM for more than three times as long as the combination of SiR-CA and Halo-TOMM20, and for 16-times longer than the fluorescent protein marker mEmerald-TOMM20. When paired with HMDS_655_-DBCO (**Fig. 1d**) and visualized using SMLM, MAO-N_3_ distinguished discrete IMM cristae structures that were not resolved using RhoB-HMSiR, a previously reported HIDE probe assembled using RhoB-TCO and HMSiR-Tz^17^. When paired with SiR-DBCO and visualized using STED, MAO-N_3_ again improved visualization of discrete IMM cristae structures relative to RhoB-SiR^20^, and supported the acquisition of a 125-frame STED movie showing cristae dynamics.

## Results

### HIDE probe design and synthesis

The design of inner mitochondrial membrane (IMM)-selective HIDE probes was inspired by the properties of nonyl acridine orange (NAO), a small molecule fluorophore that localizes selectively to the inner mitochondrial membrane (IMM)^22^. Selectivity is achieved by favorable, non-covalent interactions with cardiolipin, an unconventional lipid that is enriched within the IMM^23^. Despite its selective localization, NAO is not photostable and cannot support long time-lapse imaging, especially using modalities that demand high-intensity irradiation^22,24^. We designed two NAO analogs that could assemble HIDE probes suitable for long time-lapse imaging for the IMM: HAO-N_3_ and MAO-N_3_ (**Fig. 1c**). HAO-N_3_ was synthesized from acridine orange *via N*-alkylation (**Extended Data Fig. 1a**)^24^, while MAO-N_3_ made use of a Minisci-type radical difluoroalkylation^25^ reaction followed by methylation (**Extended Data Fig. 1b**).

### Online Methods

Plots show the relative bioluminescence signals (% relative to untreated cells). Untreated cells serve as a negative control; cells treated with 5 μM alexidine serve as a positive control. Each set of conditions include 12 biological replicates. **d,** Pearson’s colocalization coefficients (PCC) representing the colocalization of 750 nM SiR-DBCO with 100 nM MAO-N_3_ or PDHA1-GFP (a *bona fide* mitochondrial matrix marker). PCC (SiR/PDHA1-GFP) = 0.72 ± 0.01, PCC (SiR/MAO) = 0.65 ± 0.01. *n* represents the number of cells sourced from at least two biological replicates. Error bars = s.e.m. **** p<0.0001, *** p<0.001, ** p<0.01, *p<0.1, n.s. = not significant, from one-way ANOVA with Dunnett’s post-analysis accounting comparison to the negative control where cells were incubated with cell culture media only (for **c**) or Tukey’s multiple comparison test (for **d**).

### *In vitro* characterization of HAO-N_3_ and MAO-N_3_

We first evaluated the *in vitro* photophysical properties of HAO-N_3_ and MAO-N_3_ in comparison with the parent fluorophore NAO. NAO exhibits an absorption maximum at 496 nm, an emission maximum at 519 nm, and a quantum yield of 0.16^26^. The photophysical properties of HAO-N_3_ are similar, with an absorption maximum at 496 nm, and an emission maximum at 517 nm (**Fig. 2a** TOP and **Extended Data Fig. 2a-d**). In contrast, the absorption and emission curves for MAO-N_3_ were red-shifted^27^, with an absorption maximum at 520 nm and an emission maximum at 570 nm (**Fig. 2a** BOTTOM and **Extended Data Fig. 2e-h**). MAO-N_3_ (QY = 0.01, ε = 3.0 × 10^4^ L·mol^−1^·cm^−1^) is also dimmer than HAO-N_3_ (QY = 0.11, ε = 6.8 × 10^4^ L·mol^−1^·cm^−1^). The red shift and lower brightness of MAO-N_3_ are both advantageous for the lipid-like component of a HIDE probe. These properties minimize cross-talk with GFP (λ_ex_ = 490 nm, λ_em_ = 520 nm) and RFP (λ_ex_ = 555 nm, λ_em_ = 583 nm) organelle markers as well as common small molecule fluorophores that would otherwise complicate multicolor imaging experiments. *In vitro* experiments verified that an aqueous solution of MAO-N_3_ and HMDS_655_-DBCO reacted within minutes *in vitro* at 37 °C to generate the expected SPAAC product, MAO-HMDS_655_ (**Fig. 2b, Supplementary Fig. 1a**).

**Fig. 2.**
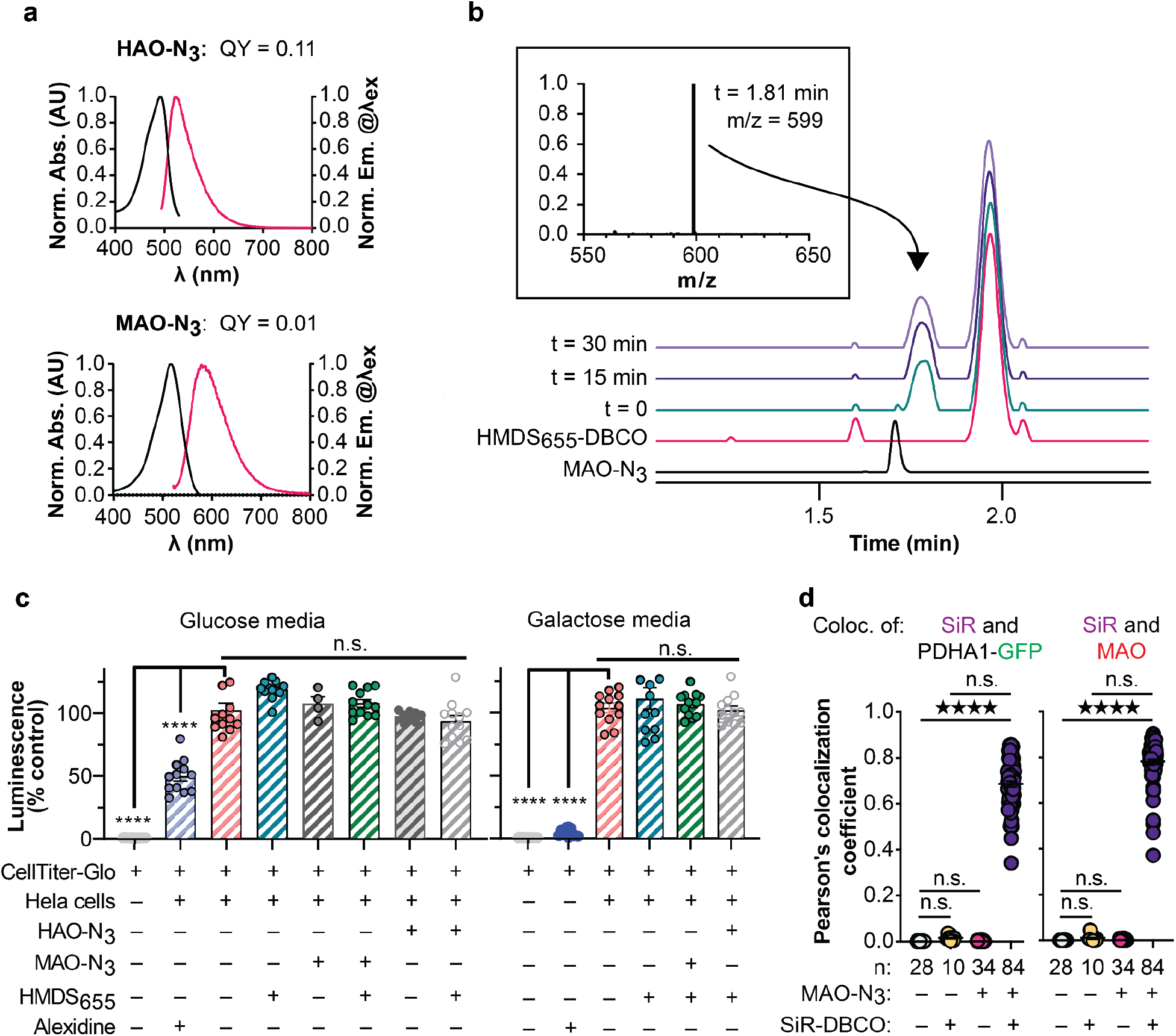
Assembly and characterization of HIDE probes generated using HAO-N_3_ and MAO-N_3_. **a,** Absorption and emission spectra and quantum yield of HAO-N_3_ and MAO-N_3_ at 2 μM concentration in Dulbecco’s phosphate-buffered saline (DPBS, pH 7.4, room temperature). **b,** *In vitro* SPAAC reaction of 100 μM MAO-N_3_ and 400 μM HMDS_655_-DBCO at 37 °C in DI H_2_O. At the times indicated, 1 μL aliquots of the reaction mixture were withdrawn and analyzed by LC/MS as described in **Online Methods**. **Insert**: Mass spectrum confirming the formation of MAO-HMDS_655_ (MW = 1197.6, m/z = 599), which elutes at 1.81 min. **c,** HeLa cells incubated in standard media (DMEM with 4.5 g/L glucose, supplemented with 10% FBS) or oxidative media (DMEM with 4.5 g/L galactose, supplemented with 10% FBS) were treated with MAO-N_3_, HAO-N_3_ and/or HMDS_655_-DBCO and the ATP levels were measured immediately as described in

### HIDE probes generated using HAO-N_3_ and MAO-N_3_ are non-toxic

Next we evaluated the effect of HAO-N_3_, MAO-N_3_, and HIDE probes derived thereof on the viability of HeLa cells under standard conditions (DMEM with 4.5 g/L glucose, supplemented with 10% FBS) and oxidative conditions (DMEM with 4.5 g/L galactose, supplemented with 10% FBS) (**Fig. 2c** and **Extended Data Fig. 3**). Cells grown in oxidative media are reliant on oxidative phosphorylation and thus sensitive to mitochondrial toxins^28^. Cell viability was evaluated using a commercial assay that detects ATP (CellTiter Glo® 2.0). As expected, cells treated with the known mitochondrial toxin alexidine^29^ displayed diminished ATP levels, and the effects were larger when cells were incubated in oxidative media containing galactose in place of glucose. The ATP levels of cells treated for 1 h with 100 nM MAO-N_3_ or HAO-N_3_, with or without co-incubation with 750 nM HMDS_655_-DBCO (**Fig. 2c**) or 750 nM SiR-DBCO (**Extended Data Fig. 3**), did not differ significantly from the ATP levels in untreated cells. We also treated cardiomyocytes differentiated from human induced pluripotent stem cells (hiPSC) with 100 nM MAO-N_3_ followed by 750 nM SiR-DBCO. No apparent effect on the rate of cell body contractions was observed over the course of 110 s (**Supplementary Movie 1**).

**Fig. 3.**
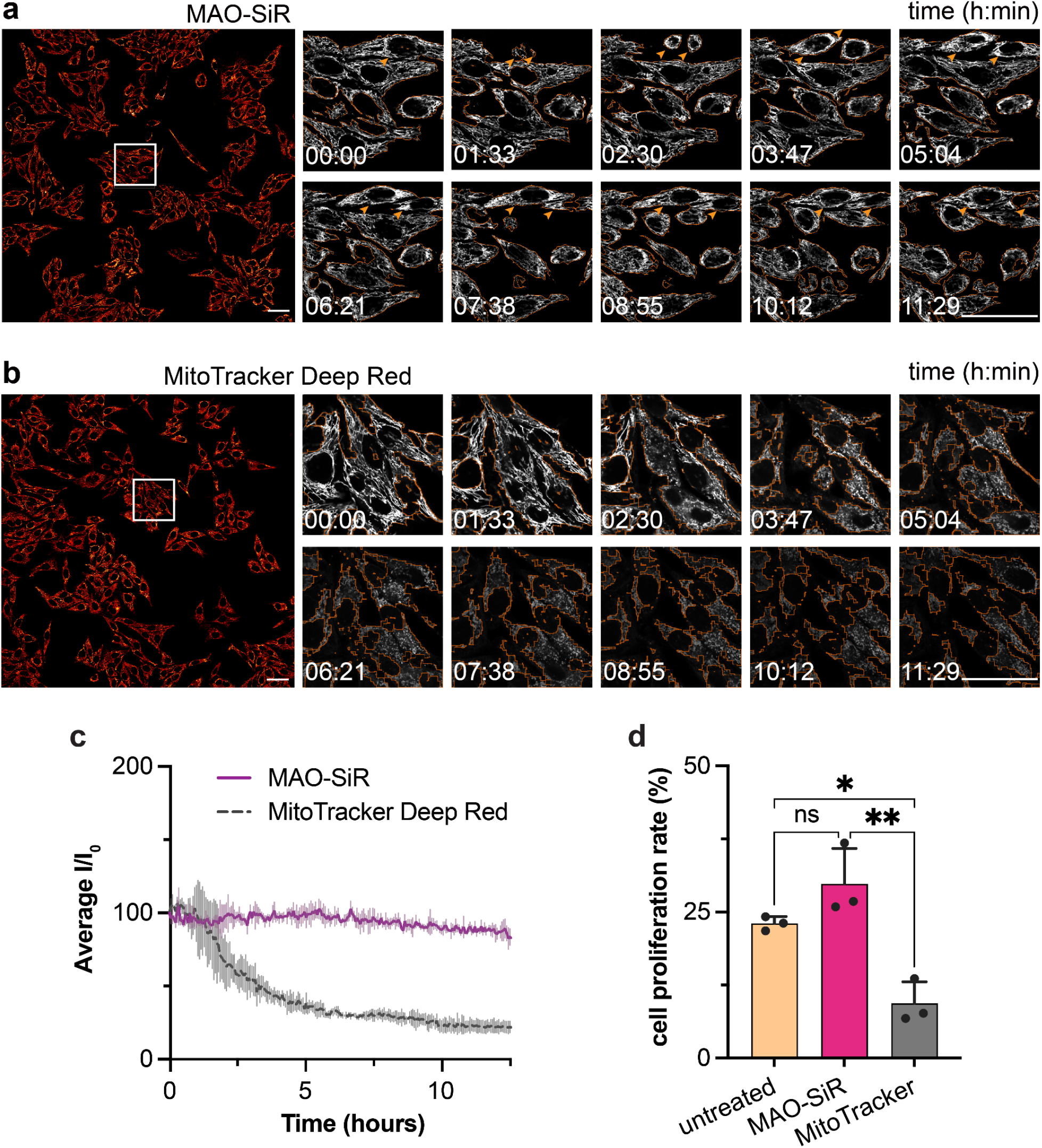
Long time-lapse confocal image using MAO-SiR or MitoTracker Deep Red. **a-b,** HeLa cells were labeled with **a**, 200 nM MAO-N_3_ and 100 nM SiR-DBCO or **b**, 100 nM MitoTracker Deep Red and imaged using a point-scanning confocal microscope. A total of 220 frames of 670 × 670 μm^2^ images were obtained over 12.5 h. Individual frames from regions of interest (white rectangle) shown to the right. Orange outlines indicate segmentation boundaries used to isolate signals for intensity quantification. Orange arrows in panel **a** represent observable cell division events. Scale bar: 50 μm. **c**, Plots of average signal decay over time. Mean signals from segmented regions were averaged to determine the mean frame intensity, Mean frame intensities from each dataset were averaged and plotted for each time point along with the relative standard deviation. *n* = 3 replicates. t_1/2_ (MitoTracker) = 1.96 ± 0.05 h, t_1/2_ (MAO-SiR) = N.A. Intensities for plots in **c** were scaled relative to the first frame’s intensity in the data sets. **d**, Cell proliferation rates for untreated, MAO-SiR or MitoTracker Deep Red labeled HeLa cells over the entire FOV over 12.5 h.

### HIDE probes derived from MAO-N_3_ and HAO-N_3_ localize to mitochondria

Confocal microscopy was used to quantify the extent to which MAO-SiR, assembled *in cellula* using MAO-N_3_ and SiR-DBCO (**Supplementary Fig. 1b**), colocalized with a *bona fide* mitochondrial marker in live HeLa cells. As a marker, we used a fusion of GFP with the leader sequence of pyruvate dehydrogenase (E1) alpha subunit-1 (PDHA1-GFP), which localizes to the mitochondrial matrix^30^ and whose emission is separate from that of MAO and SiR. Mitochondria in HeLa cells expressing PDHA1-GFP appeared as multiple discontinuous tubular structures emblematic of healthy mitochondria^2,31^ in the presence or absence of 100 nM MAO-N_3_ and/or 750 nM SiR-DBCO (**Extended Data Fig. 4b**). Cells treated with MAO-N_3_ alone showed significant signal at 570 nm, near the MAO emission maximum, which colocalized with the emission associated with PDHA1-GFP (PCC = 0.77 ± 0.01, **Extended Data Fig. 4c**). Cells treated with SiR-DBCO in the absence of MAO-N_3_ showed negligible signal at 660 nm, near the SiR emission maximum. However, cells treated with both MAO-N_3_ and SiR-DBCO showed evidence of signal from SiR, which colocalized well with that of PDHA1-GFP and MAO (PCC = 0.72 ± 0.01 and 0.65 ± 0.01, respectively) (**Fig. 2d**). Although HeLa cells treated with 100 nM NAO and 750 nM SiR-DBCO showed significant signal at 520 nm, near the NAO emission maximum, negligible signal in the SiR channel was observed–illustrating that the azido group of MAO-N_3_ and a subsequent SPAAC reaction^32,33^ is essential to recruit SiR-DBCO to the mitochondria (**Extended Data Fig. 5**). Additional experiments demonstrated minimal colocalization of MAO-SiR with either the ER or the Golgi (**Extended Data Fig. 6**), using corresponding *bona fide* markers (GALNT1-GFP^34^ for Golgi and KDEL-GFP^35^ for ER). Analogous experiments using HAO-N_3_, SiR-DBCO, and RFP-tagged organelle markers (**Extended Data Fig. 7**), confirmed that HAO-N_3_ is also capable of mitochondria-specific labeling. We conclude that HAO-N_3_ and MAO-N_3_ localize selectively to mitochondria in live HeLa cells, and formation of the corresponding HIDE probe upon SPAAC reaction with SiR-DBCO localizes SiR to mitochondria.

### HIDE probe MAO-SiR supports confocal live-cell imaging of mitochondria for more than 12 h with no loss of signal intensity

We labeled HeLa cells with 200 nM MAO-N_3_ and 100 nM SiR-DBCO and obtained a 220-frame confocal time-lapse series with a 670 × 670 μm^2^ field of view over the course of 12.5 h (**Supplementary Movie 2**). Each frame in the time series was reconstructed by stitching 16 individual frames together. Over this time period, HeLa cells labeled with MAO-SiR exhibited negligible signal decay and increased in density by 27 ± 6%, which compares well with an increase of 23 ± 1% for untreated cells (**Supplementary Movie 3**)^36^. In comparison, cells labeled with 100 nM MitoTracker Deep Red, a commercial, carbocyanine-based mitochondria dye with a similar emission profile (λ_ex_ = 644 nm, λ_em_ = 665 nm), increased in density by only 9 ± 4%, and the signal intensity decreased by 50% within 2 h (**Supplementary Movie 4**). Although previous work demonstrated HIDE probes increase the length of time over which bright images can be acquired using SMLM and STED microscopy^14,17–21^, these experiments reveal that comparable improvements are seen even during microscopy using lower intensity irradiation.

### SIM imaging of the mitochondria using SiR-DBCO and MAO-N_3_

Having confirmed using confocal microscopy that the HIDE probe MAO-SiR is non-toxic, mitochondria-specific, and supports extended-time imaging, we turned to higher-resolution methods to evaluate its intra-organelle localization. HeLa cells expressing either mEmerald-TOMM20 (OMM marker) or PDHA1-GFP (mitochondrial matrix marker) were incubated with 75 nM MAO-N_3_ and 600 nM SiR-DBCO and imaged using lattice SIM (**Fig. 4a**). Images of treated cells expressing mEmerald-TOMM20 show clear differentiation of the OMM (mEmerald) and internal mitochondrial fine structure (SiR) (**Fig. 4b,c**). Images of treated cells expressing PDHA1-GFP (mitochondrial matrix marker) also show differences in signal localization (**Fig. 4d-f**). While a profile of the signals due to PDHA1-GFP are relatively continuous, those due to MAO-SiR are not (**Fig. 4e,f**), revealing a discontinuous structure that resembles the ones seen in **Fig. 4b**. Due to the limited resolution of SIM (**Extended Data Fig. 8a-d**), the signals from the SiR channel (FWHM = 188 ± 5 nm, **Extended Data Fig. 8d**) cannot resolve the cristae topology of the IMM^12,37^. Despite this limitation, these SIM experiments provide strong evidence that the HIDE probe MAO-SiR labels neither the OMM nor the mitochondrial matrix.

**Fig. 4.**
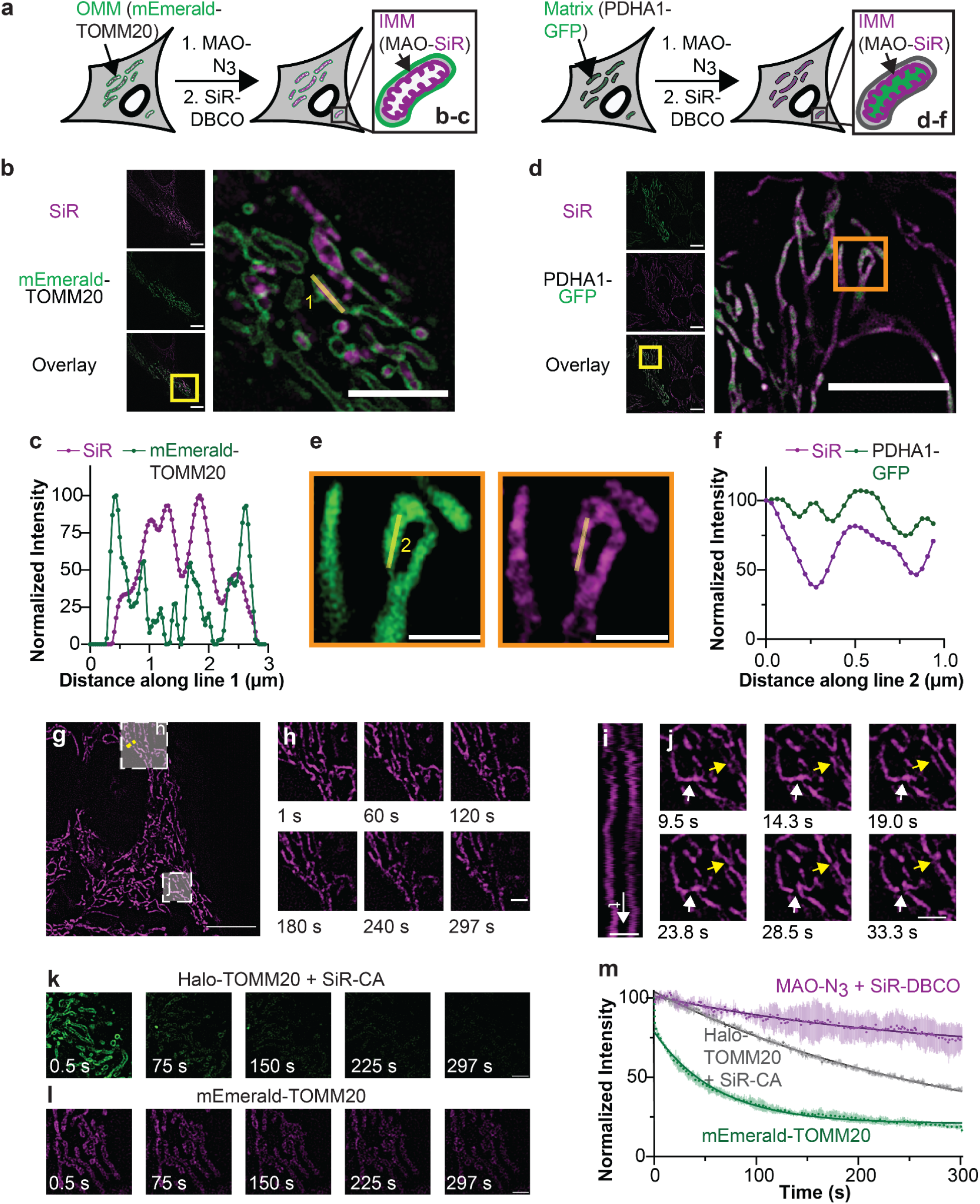
SIM imaging of mitochondria using MAO-SiR. **a,** HeLa cells expressing PDHA1-GFP or mEmerald-TOMM20 were treated with 75 nM MAO-N_3_ and 600 nM SiR-DBCO as described in **Online Methods** and imaged using lattice SIM. **b,** SIM images of cells expressing mEmerald-TOMM20 and labelled with MAO-SiR as described in **a**. Scale bar: 5 μm. **c,** Plots of signal intensity from SiR and mEmerald along line 1. **d,** SIM images of cells expressing PDHA1-GFP and labelled with MAO-SiR as described in **a**. Scale bar: 5 μm. **e,** Enlarged image within orange box in panel **d. f,** Plots of signals due to SiR and GFP along line 2. **g-j,** Long time-lapse SIM imaging of IMM dynamics using the HIDE probe MAO-SiR, Halo-TOMM20 (labeled with SiR-CA to generate SiR-TOMM20), or mEmerald-TOMM20 in HeLa cells. Panels **g-j** are representative images from a 500-frame SIM series obtained over 297 s (**Supplementary Movie 5**). **g,** SIM image at t = 0.5 s. Scale bar: 10 μm. **h,** Snapshots of the bracketed region indicated in **g** at the times indicated. Scale bar: 2 μm. **i,** Kymograph of the yellow dashed line in **g**. Scale bar: 1 μm. **j,** Mitochondria fission (white arrow) and fusion (yellow arrow) events from t = 9.5 s to t = 33.3 s in the bracketed region in **g**. Scale bar: 2 μm. **k and l,** Snapshots of selected regions of interest from **Supplementary Movie 6 and 7**, obtained from HeLa cells expressing **k,** mEmerald-TOMM20 (**Supplementary Movie 6**) or **l,** SiR-TOMM20 (**Supplementary Movie 7**), at times indicated. Scale bar: 2 μm. **m,** Plots illustrated normalized SiR fluorescence signals of SiR-MAO, SiR-TOMM20, and green fluorescent protein signal from mEmerald-TOMM20 over time. *n* = 3 regions of interest (ROI), *N* = 1 cell. Intensities for plot m were scaled relative to the first frame’s intensity in the data sets.

To evaluate the time over which interpretable images can be acquired using SIM, we treated HeLa cells with MAO-N_3_ and SiR-DBCO as described above and imaged them for 297 s (500 frames) (**Fig. 4g-j**, **Supplementary Movie 5**). Over a field of view sufficient to visualize an entire cell (**Fig. 4g**), this lattice SIM movie captured both mitochondrial fission (white arrow) and fusion (yellow arrow) events (**Fig. 4j**) with no change in mitochondrial morphology (**Fig. 4h**) and excellent signal retention (**Fig. 4i**). When cells were imaged using mEmerald-TOMM20, little or no signal was observed after 75 s (**Fig. 4k, Supplementary Movie 6**). When cells expressing Halo-TOMM20-N-10 were labeled with SiR-HaloTag-ligand (SiR-CA) to generate the fluorophore-protein conjugate SiR-TOMM20, significant signal loss was also evident (**Fig. 4l, Supplementary Movie 7**). Exponential decay analyses of intensity data revealed a signal half-life of 713.5 ± 8.0 s for MAO-SiR, which is 3.2 times as long as that of SiR-CA/Halo-TOMM20 (t_1/2_ = 224.4 ± 0.6 s) and 16 times longer than mEmerald-TOMM20 (t_1/2_ = 42.4 ± 0.5 s) (**Fig. 4jm**). The signal half-life of HAO-SiR (t_1/2_ = 248.2 ± 1.2 s) and the previously reported mitochondrial HIDE probe RhoB-SiR (t_1/2_ = 233.0 ± 1.3 s) were both also significantly smaller than MAO-SiR (**Extended Data Fig. 9**).

### SMLM imaging of the mitochondria using HMDS_655_-DBCO and MAO-N_3_

We next used an MAO-derived HIDE probe and SMLM to visualize IMM fine structure. HeLa cells were treated with 75 nM MAO-N_3_ for 40 min, washed, incubated for 20 min with 600 nM HMDS_655_-DBCO^38^ (**Fig. 5a**), and imaged using SMLM in an antioxidant-supplemented buffer^17^. A representative image constructed from 800 frames (t = 4 s) is shown in **Fig. 5b**. Although the low temporal resolution made it challenging to capture the fine structure of continuously moving mitochondria^39^, the images reveal multiple tubular cristae structures in a ladder-like pattern; these features were not resolved by a previously reported HIDE probe generated using RhoB-TCO and HMSiR-Tz^17^. The resolution of the images using MAO-HMSiR is sufficiently high (FWHMs of 76 ± 6 nm and 81 ± 7 nm for representative peaks in the line profile) (**Fig. 5c**) to establish distances between adjacent cristae of 68 nm to 184 nm, comparable to previous reports^12,37^.

**Fig. 5.**
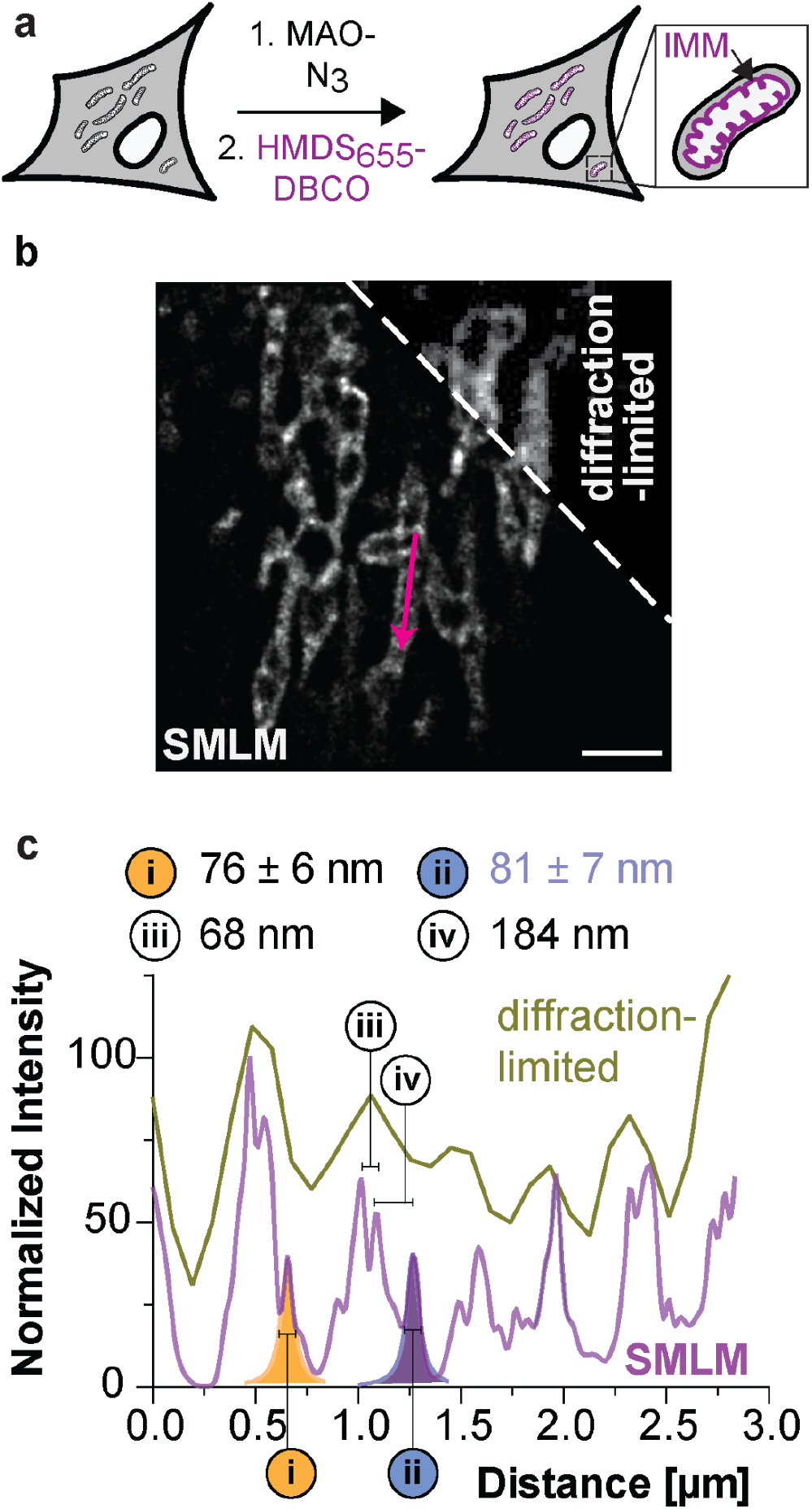
SMLM imaging of the inner mitochondrial membrane using a HIDE probe assembled from MAO-N_3_ and HMDS_655_-DBCO (MAO-HMDS_655_). **a,** HeLa cells were treated with 75 nM MAO-N_3_ and 600 nM HMDS_655_-DBCO and imaged using a widefield microscope with SMLM capability. **b,** Representative SMLM image reconstructed with 800 frames (t = 4 s); the corresponding diffraction-limited image is shown for comparison. Scale bar: 2 μm. **c,** Plot of signal intensity along the magenta arrow in panel **b** in both the reconstructed SMLM and wide-field images. Two peaks in the SMLM image labeled **i** and **ii** were fitted into a Lorentzian function to obtain FWHM values of 76 ± 6 nm and 81 ± 7 nm, respectively. Distances (**iii** and **iv**) between two pairs of adjacent peaks were measured from centroid to centroid, giving values of 68 nm and 184 nm, respectively.

Scale bar: 2 μm. **c**, Plots illustrated relative SiR fluorescence signals of SiR-MAO and SiR-CA/Halo-TOMM20, over 125 frames obtained in 162.5 s. *N* = 7 FOVs over 7 individual cells. Intensities for plot **c** were scaled relative to the first frame’s intensity in the data sets. **d-e**, Selected STED images using MAO-SiR (panel **d**, yellow, **Supplementary Movie 8**) or SiR-CA/Halo-TOMM20 (panel **e**, blue, **Supplementary Movie 9**), each from a 125-frame, 162.5 s movie at indicated time points. White bar in each frame indicates the region of interest (ROI) for the kymograph in the right most panel. Kymograph plotted to the right of each frame set to track intensity of signal in ROI for each frame in the movie. Scale bar: 1 μm.

### STED imaging of IMM cristae enabled by SiR-DBCO and MAO-N_3_

Having validated that HIDE probes assembled using MAO-N_3_ offer tangible improvements in the time over which IMM images can be generated using both SIM and SMLM, we turned to the toughest test - STED. Previous IMM markers that support STED imaging, such as Cox8A-SNAP-SiR^40^, MitoPB Yellow^12^, PKMO^13^, and MitoEsq635^8^, are limited by photostability or photocytotoxicity.

We first examined different data acquisition modes in STED microscopy, using HeLa cells labeled with MAO-SiR. We acquired images of the cristae using gated STED and deconvolved the images using a theoretical point spread function (PSF) (FWHM up to 63 ± 2 nm, **Fig. 6a**, TOP). We then filtered the fluorescence signals on the basis of fluorescence lifetime in tauSTED mode^41^ and acquired images of the IMM with excellent resolution (**Fig. 6a**, BOTTOM). These images clearly demark densely-packed, ladder-like mitochondrial cristae with regular dimensions (**Fig. 6b**) that are indiscernible using confocal microscopy (**Extended Data Fig. 10 a,b**). Moreover, the fluorescent signal from MAO-SiR diminshed 10-times slower (half-life = 112.7 ± 2.1 s) than signal from SiR-TOMM20 (half-life = 10.4 ± 0.2 s) (**Fig. 6c**). This apparent photostability supported the acquisition of a 162.5 s movie of cristae dynamics containing 125 tauSTED frames (**Fig. 6d-e**). Distinct cristae are visible in images obtained with MAO-SiR until the last frame without the need for genetic manipulation (**Fig. 6d, Supplementary Movie 8**), whereas images obtained with SiR-TOMM20 are lost within 65 s (**Fig. 6e, Supplementary Movie 9**).

**Fig. 6.**
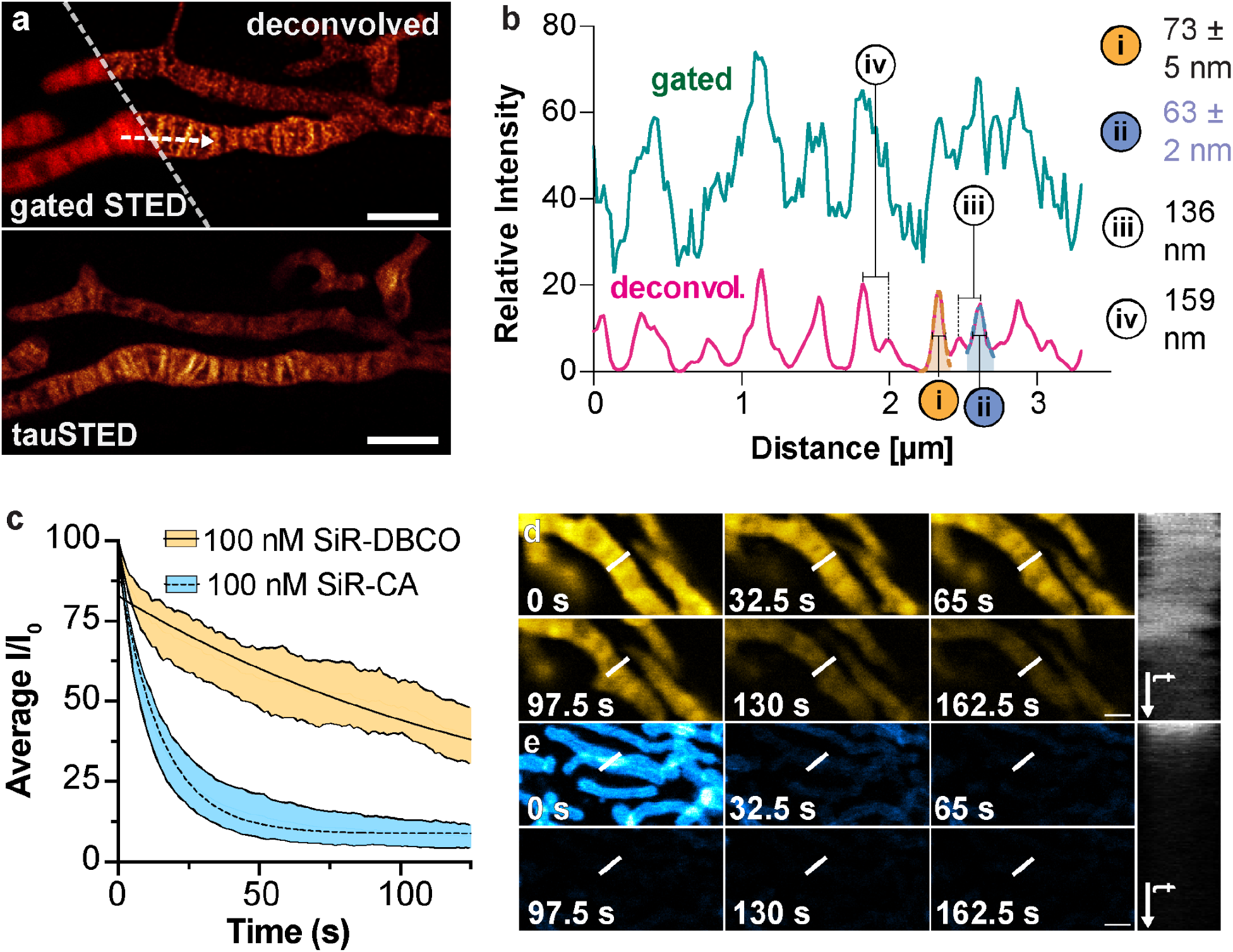
STED imaging of mitochondrial IMM using MAO-SiR. **a**, Top: gated and deconvolved STED image; Bottom: tauSTED image generated using fluorescence lifetime filtering. **b**, Plot of SiR signal intensity along white arrow in **a** in the gated STED and deconvolved STED image. Two peaks in the STED image were fitted into a Lorentzian function to obtain an FWHM of (**i**) 73 ± 5 nm and (**ii**) 63 ± 2 nm, respectively. Distances between pairs of adjacent peaks were also measured from centroid to centroid, giving values of (**iii**) 136 nm and (**iv**) 159 nm, respectively.

## Discussion

The mitochondria is a highly dynamic organelle that undergoes coordinated and balanced cycles of fission and fusion to maintain its shape, content, transport ability, and size. These transient and rapid morphological changes allow mitochondria to meet metabolic demands, initiate the degradation of damaged organelles, and contribute to control of the cell cycle, immunity, and cellular apoptosis. Defects in mitochondrial dynamics cause disease^42^. Decoupled fusion and fission and the resulting mitochondrial fragmentation occur in response to stress and during heart failure^43^, neurodegenerative disease, cancer, and obesity. Recent studies have shown that the dynamics of the inner mitochondrial membrane are critical, for example, for the loss of mitochondrial DNA (mtDNA) from the matrix into the cytosol to trigger the innate immune cGAS/STING pathway^44^.

Imaging the mitochondria to detect dynamic morphological changes–especially those involving the complex inner membrane–is especially challenging, as the subcompartment is replete with enzymes that generate dye-inactivating reactive oxygen species. Moreover, the imaging process itself can damage essential macromolecules. These challenges are exacerbated when one seeks to evaluate changes in mitochondrial dynamics or interactions that occur over significant time frames, in primary or patient-derived cells, and/or using super-resolution imaging modalities that require intense irradiation. One proven strategy to overcome these challenges involves labeling the organelle membrane lipids as opposed to the proteins embedded within^14–21,45^. Labeling the membrane lipids positions the dye at high density within a hydrophobic environment in which most dye molecules exist in a dark state and do not absorb light. Balancing the labeling density with the fraction of molecules in the dark state produces bright images that last because the dark state molecules act as a reservoir to replenish dyes that bleach.

Here we describe new HIDE probes that selectively image an organelle sub-compartment – the inner mitochondrial membrane – in live cells and over exceptional long time frames. These probes, all assembled from the lipid-like molecule MAO-N_3_, are photostable, lack detectable toxicity, and support exceptional spatial resolution of the IMM using multiple modalities, all without genetic manipulations. Confocal images of the mitochondria generated using MAO-SiR remained stable for more than 12.5 h with little or no loss in signal intensity or cell viability, more than 6-times longer than possible using MitoTracker Deep Red. SIM images of the inner mitochondrial membrane using MAO-SiR lasted for 16-times longer than those obtained using mEmerald-TOMM20, and supported acquisition of 500 frames over the course of 5 min as opposed to 20 frames over 2.8 min possible with MitoTracker Green^46^. When paired with HMDS_655_-DBCO (**Fig. 1d**) and visualized using SMLM, MAO-N_3_ distinguished discrete IMM cristae structures that were not resolved using a previously reported HIDE probe assembled using RhoB-TCO^17^. When paired with SiR-DBCO and visualized using STED (**Fig. 1d**), MAO-N_3_ again improved visualization of discrete IMM cristae structures relative to RhoB-TCO^20^, and supported the acquisition of a 125-frame STED movie showing cristae dynamics. HIDE probes based on MAO-N_3_ are versatile, non-toxic, photostable, and completely cell-permeant. These features should make them highly useful, alone or in combination with other organelle markers, for long-term analysis of natural and pathological mechanisms related to IMM dynamics and inter-organelle interactions.

## Online Methods

### Materials

Methods used to synthesize MAO-N_3_, HAO-N_3_, HMDS_655_-DBCO and full characterization data can be found in the Supplementary Note. SiR-CA was synthesized according to previously reported methods. BacMam^®^ 2.0 reagent, CellLight Mitochondria-GFP (PDHA1-GFP) (C10600), CellLight Mitochondria-RFP (PDHA1-RFP) (C10505), CellLight Golgi-GFP (GALNT1-GFP) (C10592), CellLight Golgi-RFP (GALNT1-RFP) (C10593), CellLight ER-GFP (KDEL-GFP) (C10590), CellLight ER-RFP (KDEL-RFP) (C10591), culturing media Dulbecco’s Modified Eagle medium (DMEM, 12430054, 21063029, and A1443001), Opti-MEM with no phenol red (11058021), and fetal bovine serum (FBS), and chemical reagents Hoechst33342 (62249), MitoTracker Deep Red FM (M22426) were all purchased from ThermoFisher. CellTiter-Glo^®^ 2.0 (G9242) was purchased from Promega, and alexidine dihydrochloride (A8986) was purchased from Sigma Aldrich. Plasmid encoding mEmerald-TOMM20 was a gift from Michael Davidson (mEmerald-TOMM20-N-10, Addgene plasmid # 54282). Plasmid encoding Halo-TOMM20 was a gift from Kevin McGowan (Halo-TOMM20-N-10, Addgene plasmid # 123284).

### Cell culture

HeLa cells (UC Berkeley Cell Culture Facility) were cultured in DMEM (ThermoFisher) supplemented with 10% FBS, penicillin (100 units/mL) and streptomycin (100 μg/mL). All cells were cultured at 37 °C in a humidified CO_2_/air (5/95%) incubator. All cells were bona fide lines obtained from UCB-CCF and periodically tested for mycoplasma with DNA methods. Cells for imaging were seeded in four- or eight-well Lab-Tek chambers (Nunc, Thermo Fisher #177399 and 177402, No. 1.0, 0.8 cm^2^ for four-well and 1.8 cm^2^ for eight-well. Used for confocal colocalization studies, SIM and SMLM) or four-well glass bottom μslides (Ibidi #80427, No. 1.5H, 2.5 cm^2^, for STED and overnight confocal imaging) at indicated density.

### Microscopy

Colocalization studies was performed on a Zeiss LSM 880 confocal microscope equipped with a Plan-Apochromat ×63/1.4 NA (Zeiss) oil objective and a diode 405 nm laser, an Argon 458, 488, 514 nm laser, a diode pumped solid-state 561 nm laser, and a 633 nm HeNe laser with standard settings. The pinhole size was set to 1 airy unit.

Overnight time-lapse confocal and STED microscopy were performed on a Leica STELLARIS 8 microscope (Leica Microsystems) equipped with a Leica DMi8 CS scanhead; a HC Plan-Apo 100×/1.4NA STED White oil immersion objective (for STED), a HC Plan-Apo 63×/1.4NA oil immersion objective (for point-scanning confocal time-lapse), a HC Plan-Apo 63×/1.4NA water immersion objective (for point-scanning confocal hiPSC-CM imaging), a pulsed white-light laser (440 nm- 790 nm; 440 nm: > 1.1 mW; 488 nm: > 1.6 mW; 560 nm: > 2.0 mW; 630 nm: > 2.6 mW; 790 nm: > 3.5 mW, 78 MHz), and a pulsed 775nm STED laser. Confocal imaging was carried out using either HyD S or HyD X detectors in analog or digital mode, while STED was exclusively carried out on HyD X detectors in photon counting mode. Live cell imaging conditions were maintained using a blacked out cage enclosure from Okolab. Temperature was maintained by heating the enclosure and monitored using Oko-Touch. pH was maintained by supplying humidified 5% CO_2_ to the sample chamber.

Lattice SIM and SMLM imaging was performed on an inverted Zeiss Elyra 7 microscope equipped with a Plan-Apo 63×/1.46 NA Oil immersion objective (Zeiss) and a 403 nm (200 mW), a Sapphire 488 nm (1 W), 561 nm (1.3W) and a Lasos 642 nm (550 mW) laser, a MBS 405/488/561/641 and EF LBF 405/488/561/641 filter set, a LP 560 and a BP 570-620 + LP 655 filter cube, and two p.co edge high speed sCMOS cameras, using Zeiss Zen Black software.

### In vitro spectroscopic characterization

Solutions used for absorption and emission spectra and quantum yield were obtained by diluting a 2 mM DMSO stock solution of each fluorophore with Dulbecco’s phosphate-buffered saline (DPBS, pH 7.4) to the desired concentration (2 μM to 125 nM). Absorption spectra were measured at room temperature using a Perkin-Elmer Lambda365 spectrometer in 1 cm pathlength, 1 mL cuvettes. Extinction coefficients were determined by performing a linear fitting of absorbance maxima (490 nm for HAO-N_3_, 520 nm for MAO-N_3_) against concentration of MAO-N_3_ or HAO-N_3_. Fluorescence spectra were measured at room temperature using a Jobin Yvon Fluoromax fluorimeter using 1 cm pathlength, 3.5 mL cuvettes. Absolute quantum yield was measured on Hamamatsu Quantaurus-QY (model No. C13534) using 1 cm pathlength, 3.5 mL cuvettes.

### In vitro SPAAC reaction

In a 300 μL LCMS vial (equipped with high recovery insert and pre-slit rubber septum), 100 μL MAO-N_3_ (100 μM) and 100 μL of HMDS_655_-DBCO (400 μM) solutions in DI H_2_O were mixed and incubated at 37 °C. As is indicated in **Fig. 2d**, 1 μL aliquots of the reaction mixture were withdrawn upon mixing, after 15 min and 30 min of incubation, respectively. The aliquots were analyzed by a Waters Acquity SQD2 UPLC-MS system equipped with a 2.1 mm × 30 mm, CSH C18 column with eluents A (H_2_O with 0.1% HCO_2_H) and B (CH_3_CN with 0.1% HCO_2_H).

### Media exchange for HeLa cells in galactose supplemented media

Five days before the experiment, HeLa cells were seeded in a T-75 tissue culture treated flask at a density of 1.0 × 10^6^ cells and incubated in DMEM supplemented with 10% FBS and 4.5 g/L glucose at 37 °C, 5% CO_2_ for 24 h. On day 2, the media was exchanged with DMEM supplemented with 10% FBS, 2.25 g/L glucose and 2.25 g/L galactose and incubated for another 24h. On day 3, the media was exchanged to DMEM, 10% FBS, 1.125 g/L glucose and 3.375 g/L galactose and to media supplemented with 4.5 g/L galactose on day 4. 24 h after the final media exchange step, the HeLa cells were plated for cell viability test.

### Cell viability assays

HeLa cells were plated into the wells of a flat-bottom 96-well plate (2 × 10^4^ cells per well in 300 μL of media) and each well was incubated overnight. On the following day, the cells in each well were treated with 300 μL DMEM (negative control) or DMEM solutions of 100 nM HAO-N_3_, MAO-N_3_, or 5 μM alexidine (cytotoxicity positive control) for 1 h. Each well was then washed twice with 300 μL DMEM ph(-), followed by a 1 h incubation with 300 μL of DMEM pH(-) solutions of either 750 nM HMDS_655_-DBCO or 750 nM SiR-DBCO. The media in each well was then exchanged with 300 μL DMEM supplemented with 10% FBS and incubation continued for 20 min. The buffer exchange and incubation was repeated for a total of 3 times. The media in each well was discarded, replaced with 100 μL of DPBS, and 100 μL CellTiterGlo 2.0 (Promega) solution was added. The plate was incubated at room temperature for 15 min in the dark. The bioluminescent level of each well was measured using a Synergy H1 plate reader (96-well opaque-walled tissue culture plate, 22 °C). Each of the relative bioluminescence signals (% relative to untreated cells) shown represents the average of 12 replicates.

### Confocal imaging to monitor cell body contraction of human induced pluripotent stem cell-derived cardiomyocytes (hiPSC-CM)

hiPSC-CM were prepared according to reported procedure^47^, and cultured on Matrigel (1:100 dilution; Corning)-coated 24 well μ-plates (Ibidi^®^, cat no. 82426) (7.0 × 10^4^ cells per well) in RPMI 1640 medium (Life Technologies) with B27 supplement (Life Technologies). On the day of imaging, the hiPSC-CMs were labeled with 600 μL of 100 nM MAO-N_3_ in RPMI 1640/B27 ph(-) for 1 h, washed (3 × 600 μLRPMI 1640/B27 ph(-)), incubated with 600 μL of 0.75 μM SiR-DBCO in RPMI 1640/B27 ph(-) for 1 h, and washed again (RPMI 1640/B27 ph(-), 6 × 10 min). Finally, the cells were imaged in 700 μL RPMI 1640/B27 ph(-) at 37°C and 5% CO_2_. A total of 213 frames from a field of view of 72.5 μm × 72.5 μm was obtained with a time interval of 0.52 s, with a time-lapse of 110 s. Pixel size: 0.142 μm, pixel dwell time: 0.95 μs, scan speed: 1000 Hz, laser power: 25.7 % (652 nm).

### Confocal imaging for colocalization with organelle markers

Two days before imaging, HeLa cells were seeded in LabTek-I eight-well chambers (1.5 × 10^4^ cells per well). After 24 hours, 6 μL of a CellLight BacMam reagent (40 particles per cell) for organelles of interest (ER, Golgi or mitochondria) was added to each well and incubation was continued for 16–20 h. The cells were subsequently treated with 300 μL of 100 nM HAO-N_3_ or MAO-N_3_ (diluted from 0.2 mM DMSO stock solution into DMEM ph(-) buffer) for 1 hour. The cells were washed 3× with DPBS and treated with 300 μL of a 750 nM solution of SiR-DBCO in DMEM ph(-) for 1 h. Each well was then washed with 300 μL DMEM supplemented with 10% FBS and incubated for 10 min. The wash/incubation was repeated 6 times to minimize non-specific dye labeling. Cells were then incubated with 300 μL of 0.5 μg/mL Hoechst33342 (ThermoFisher, #62249) solution for 1 min to label nuclei and help identify the plane of view. Cells were washed with warm DPBS and imaged in DMEM ph(-) at room temperature Pairwise co-localization of HAO/MAO/SiR/GFP markers was evaluated by measuring the overlap of fluorescent signals from two spectrally separated channels using the PCC for individual cells using the JaCOP plugin on the FIJI software. PCC values for each condition obtained from multiple cells collected over at least 2 biological replicates were pooled and represented as Mean with S.E.M using Prism 8.4.3.

### Overnight confocal imaging for cell proliferation monitoring

48 hours before imaging, HeLa cells (P3-P10) were seeded in a four-well chamber (2.2 × 10^4^ cells per well). On the day of the experiment, the cells were treated with 700 μL of a 200 nM solution of MAO-N_3_ (diluted from 0.2 mM DMSO stock solution in OptiMEM ph(-) buffer) for 1 hour. The cells were washed 1× with DMEM ph (−) and treated with 700 μL of a 100 nM solution of SiR-DBCO (diluted from 2 mM DMSO stock solution into OptiMEM ph(-) buffer) for 1 h. The cells were then washed with 700 μL DMEM supplemented with 10% FBS and incubated in the same buffer for 1h. For experiment with MitoTracker Deep Red, the cells were treated with 700 μL of a 100 nM solution of MitoTracker Deep Red (diluted from 0.2 mM DMSO stock solution into OptiMEM ph(-) buffer) for 30 min, and washed 3× with 700 μL of DPBS. For untreated cells, the cells were washed 3× with 700 μL of DPBS. Finally, the cells were imaged in 700 μL DMEM ph(-) with 10% FBS, at 37°C and 5% CO_2_.

A region of 4 × 4 frames was imaged every 2 minutes, and each time-point in the time-lapse was derived by stitching the 16 frames together. Each time-point captured a field of view of 670 μm × 670 μm. For each frame within the time-point composite, the field of view was 184.52 × 184.52 μm. Focal plane was maintained using Leica’s Adaptive Focus Control system. Frame size: 1024 × 1024, pixel size: 0.18 μm, pixel dwell time: 2.8375 μs, detection window: 662 - 749 nm. laser power: 10% (652 nm) for MAO-SiR and untreated cells, 2% for MitoTracker DeepRed-treated cells. Cell counts of frame #1 and frame #220 were performed manually using the Cell Counter plugin in FIJI. Segmentation was run to isolate cell bodies from background, and a mask was generated to identify ROIs for intensity quantification. Mean signal for each ROI was averaged for one time point to give the mean frame intensity. Each mean frame intensity was scaled relative to the first frame of the time series. Then the mean for each frame was averaged over three data sets to determine the half-life. Half-life of signal intensity was obtained via performing a nonlinear regression of the exponential decay function using OriginLab OriginPro 2022b.

### Two color SIM imaging of OMM and IMM using mEmerald-TOMM20 and MAO-SiR

Two days before imaging, HeLa cells were plated (2.5 × 10^4^ cells per well) in a 4-well chamber and incubated overnight. After 16 hours, the cells were transfected with plasmids encoding mEmerald-TOMM20 using FuGENE HD transfection reagent (Promega) according to manufacturer’s protocol, and incubated overnight. On the following day, the HeLa cells were labeled with 600 μL of 75 nM MAO-N_3_ in DMEM ph(-) for 1 h, washed (3 × 600 μL DMEM ph(-)), incubated with 600 μL of 0.6 μM SiR-DBCO in DMEM ph(-) for 1 h, and washed again (DMEM ph(-) 10% FBS, 6 × 10 min). The sample was subsequently visualized via Lattice SIM imaging on a Zeiss Elyra 7 microscope. Laser power: 6% (642 nm, estimated at 0.04 kw/cm^2^) and 3% (488 nm, estimated at 0.04 kw/cm^2^), field of view: 80.14 μm × 80.14 μm, camera exposure time: 40 ms, 13 phases were acquired for SIM reconstruction. Beam splitter: LBF 405/488/561/642, filter set: BP 495-550 + LP 655. Incubation conditions: DMEM ph(-) with 10% FBS, 37°C, 5% CO_2_. SIM reconstruction was performed on the Zeiss Zen Black software. Nonlinear regression for Gaussian and Lorentzian fitting of peaks from line profiles was performed using OriginLab OriginPro 2022b and plotted using Graphpad Prism 8.4.3.

### Two color SIM imaging of mitochondrial matrix and IMM using BacMam 2.0 MitoGFP and MAO-SiR

Two days before imaging, HeLa cells were plated (2.5 × 10^4^ cells per well) in a 4-well chamber and incubated overnight. After 16 hours, the cells were transfected with 10 μL Celllight BacMam 2.0 MitoGFP for 8 hrs. The cells were then washed (3×600 μL DMEM ph(-) with 10% FBS), and incubated overnight. On the following day, the HeLa cells were labeled with 600 μL of 75 nM HAO-N_3_ in DMEM ph(-) for 1 h at 37 °C with 5% CO_2_, washed (3× 600 μL DMEM ph(-)), incubated with 600 μL of 0.6 μM SiR-DBCO in DMEM ph(-) for 1 h at 37 °C with 5% CO_2_, and washed again (DMEM ph(-) 10% FBS) for 6×10 min. The sample was subsequently visualized via Lattice SIM imaging. Laser power: 6% (642 nm, estimated at 0.04 kw/cm^2^) and 3% (488 nm, estimated at 0.04 kw/cm^2^), field of view: 80.14 μm × 80.14 μm, camera exposure time: 40 ms, 13 phases were acquired for SIM reconstruction. Beam splitter: LBF 405/488/561/642, filter set: BP 495-550 + LP 655. Incubation conditions: DMEM ph(-) with 10% FBS, 37°C, 5% CO_2_. SIM reconstruction was performed on the Zeiss Zen Black software. Nonlinear regression for Gaussian and Lorentzian fitting of peaks from line profiles was performed using OriginLab OriginPro 2022b and plotted using Graphpad Prism 8.4.3.

### Time-lapse SIM imaging of IMM using MAO-SiR

HeLa cells were plated (3.0 × 10^4^ cells per well) in 4-well LabTekII chambers (Nunc, Thermo Fisher Scientific) and incubated overnight. On the following day, the HeLa cells were labeled with 600 μL of 75 nM HAO-N_3_ in DMEM ph(-) for 1 h at 37 °C, washed (3× 600 μL DMEM ph(-)), incubated with 600 μL of 0.6 μM SiR-DBCO in DMEM ph(-) for 1 h at 37 °C with 5% CO_2_, and washed again (DMEM ph(-) 10% FBS) for 6×10 min. The time-lapse SIM imaging of labeled cells was performed on a Zeiss Elyra 7 microscope using Zeiss Zen Black software. Laser power: 0.4 % (642 nm, estimated at 0.0028 kw/cm^2^), field of view: 80.14 μm × 80.14 μm, camera exposure time: 40 ms, 13 phases per frame were acquired for SIM reconstruction. 100 ms interval between each frame, 500 frames in total over an acquisition time of 297 s. Beam splitter: LBF 405/488/561/642; filter set: SBS LP 560. SIM reconstruction was performed on the Zeiss Zen Black software. Imaging buffer and conditions: DMEM ph(-) with 10% FBS, 37°C, 5% CO_2_). Kymograph of line profile (**Fig. 4g**) was obtained using the KymographBuilder plugin on the FIJI software. Fluoresecence intensity was obtained from randomly picked 3 μm × 3 μm regions using ImageJ and plotted using Graphpad Prism 8.4.3. Half-life of signal intensity was obtained via performing a nonlinear regression of the exponential decay function using OriginLab OriginPro 2022b.

### Time-lapse SIM imaging of OMM using mEmerald-TOMM20

Two days before imaging, HeLa cells were plated (2.5 × 10^4^ cells per well) in a 4-well chamber and incubated overnight. After 16 hours, the cells were transfected with plasmids encoding mEmerald-TOMM20 using FuGENE HD transfection reagent (Promega) according to manufacturer’s protocol, and incubated overnight. On the following day, the HeLa cells were visualized via Lattice SIM imaging on a Zeiss Elyra 7 microscope. Laser power: 0.4 % (488 nm, estimated at 0.005 kw/cm^2^), field of view: 80.14 μm × 80.14 μm, camera exposure time: 40 ms, 13 phases per frame were acquired for SIM reconstruction. 100 ms interval between each frame, 500 frames in total over an acquisition time of 297 s. Beam splitter: LBF 405/488/561/642; filter set: SBS LP 560. Imaging buffer and conditions: DMEM ph(-) with 10% FBS, 37°C, 5% CO_2_). SIM reconstruction was performed on the Zeiss Zen Black software. Fluorescence intensity was obtained from randomly picked 3 μm × 3 μm regions using ImageJ and plotted using Graphpad Prism 8.4.3. Half-life of signal intensity was obtained via performing a nonlinear regression of the exponential decay function using OriginLab OriginPro 2022b.

### Time-lapse SIM imaging of OMM using Halo-TOMM20 and SiR-CA

Two days before imaging, HeLa cells were plated (2.5 × 10^4^ cells per well) in a 4-well chamber and incubated overnight. After 16 hours, the cells were transfected with plasmids encoding Halo-TOMM20 using FuGENE HD transfection reagent (Promega) according to manufacturer’s protocol, and incubated overnight. On the following day, the HeLa cells were incubated with 600 μL of 2 μM SiR-CA in DMEM ph(-) for 1 h at 37 °C with 5% CO_2_, and washed 3× (600 μL of DMEM pH(-) supplemented with 10% FBS). visualized via Lattice SIM imaging on a Zeiss Elyra 7 microscope. Laser power: 0.4 % (488 nm, estimated at 0.005 kw/cm^2^), field of view: 80.14 μm × 80.14 μm, camera exposure time: 40 ms, 13 phases per frame were acquired for SIM reconstruction. 100 ms interval between each frame, 500 frames in total over an acquisition time of 297 s. Beam splitter: LBF 405/488/561/642; filter set: SBS LP 560. Imaging buffer and conditions: DMEM ph(-) with 10% FBS, 37°C, 5% CO_2_. SIM reconstruction was performed on the Zeiss Zen Black software. Fluorescence intensity was obtained from randomly picked 3 μm × 3 μm regions using ImageJ and plotted using Graphpad Prism 8.4.3. Half-life of signal intensity was obtained via performing a nonlinear regression of the exponential decay function using OriginLab OriginPro 2022b.

### SMLM imaging of IMM structure using HMDS_655_-DBCO and MAO-N_3_

Prior to seeding, the 4-well Lab-Tek® II (Nunc) chamber was treated by sonication for 15 min in 1M KOH, rinsed three times with deionized H_2_O, sterilized with 100% EtOH, and air dried overnight in a biological safety cabinet. HeLa cells were seeded onto pre-treated plates at a density of 3.0 × 10^4^ cells per well and incubated overnight at 37 °C with 5% CO_2_. On the following day, the cells were labeled with 600 μL of 75 nM MAO-N_3_ in DMEM ph(-) for 40 min at 37 °C with 5% CO_2_, washed (3× 600 μL DMEM ph(-)), incubated with 600 μL of 0.6 μM HMDS_655_-DBCO in DMEM ph(-) for 20 min at 37 °C with 5% CO_2_, and washed again (DMEM ph(-) with 10% FBS, 6×10 min, 37 °C with 5% CO_2_). Finally, the media was exchanged with 600 μL of DMEM pH(-) supplemented with 1% FBS, 100 μM Trolox and 500 μM sodium ascorbate. The sample was subsequently visualized via SMLM imaging on an inverted Zeiss Elyra 7 microscope equipped with a Plan-Apo 63×/1.4NA Oil immersion lens (40 % laser power for 642 nm laser, 60.57° TIRF mirror angle, TIRF-uHP mode, 2.5 ms exposure time for each frame, 6000 frames obtained in 36 s).

### STED imaging of cristae structures and their dynamics using SiR-DBCO and MAO-N_3_

HeLa cells were plated (2.2 × 10^4^ cells per well) in a 4-well #1.5 glass bottom chamber (Ibidi, #80827) and incubated for 42 h. On the day of imaging, the cells were labeled with 700 μL of 200 nM MAO-N_3_ in Opti-MEM ph(-) for 1 h, washed (1 × 700 μL DMEM ph(-)), incubated with 700 μL of 100 nM SiR-DBCO in Opti-MEM ph(-) for 1 h, washed with 700 μL DMEM supplemented with 10% FBS) and incubated for 1h. Finally the media was exchanged with DMEM ph(-), and the sample was subsequently visualized via STED imaging (Imaging conditions: 20% excitation laser power, depleted by 24% STED laser power, DMEM ph(-), 37°C, 5% CO_2_). Deconvolution of gated STED images was performed based on the theoretical point spread function (PSF) and the classic maximum likelihood estimation (CMLE)^48^ method using a commercial software package SVI Huygens Pro. Imaging parameters for time-lapse: 256 × 128, 1.3 s time interval, total of 125 frames, scan rate 100 Hz, no line accumulation/average, pixel size: 30 nm.

## Acknowledgements

This work was supported by the NIH (Grant No. 1R35GM134963 to A.S.). Work at the Molecular Foundry (LBNL) for SIM, SMLM imaging and measurement of absolute quantum yields was supported by the Office of Science, Office of Basic Energy Sciences, of the U.S. Department of Energy under Contract No. DE-AC02-05CH11231. We thank Dr. Felix Rivera-Molina (Yale) for helpful discussions on imaging conditions and Dr. Behzad Rad (Molecular Foundry at LBNL) for assists on SMLM and SIM imaging. The two-color confocal microscopy experiments to determine MAO-N_3_ and HAO-N_3_ localization were performed in the Biological Imaging Facility (BIF) of UC Berkeley. Dr. Hasan Celik and UC Berkeley’s NMR facility in the College of Chemistry (CoC-NMR) was acknowledged for spectroscopic assistance. Instruments in the CoC-NMR are supported in part by NIH S10OD024998.

## Author Contributions

S.Z. and A.S. conceived the project; S.Z., N.D., D.M., and A.S. designed experiments. S.Z. designed and synthesized MAO-N_3_ and HAO-N_3_; S.Z. and L.L synthesized DBCO-conjugated fluorophores. S.Z., N.D., and D.M. prepared HeLa samples for microscopy and performed imaging experiments. K.M., S.Z. and E.M. prepared iPSC-CM samples for microscopy. S.Z., N.D., and D.M. analyzed the data with inputs from A.S. S.Z., N.D., L.L., and A.S. wrote the manuscript.

---

**Extended Data Fig. 1.**
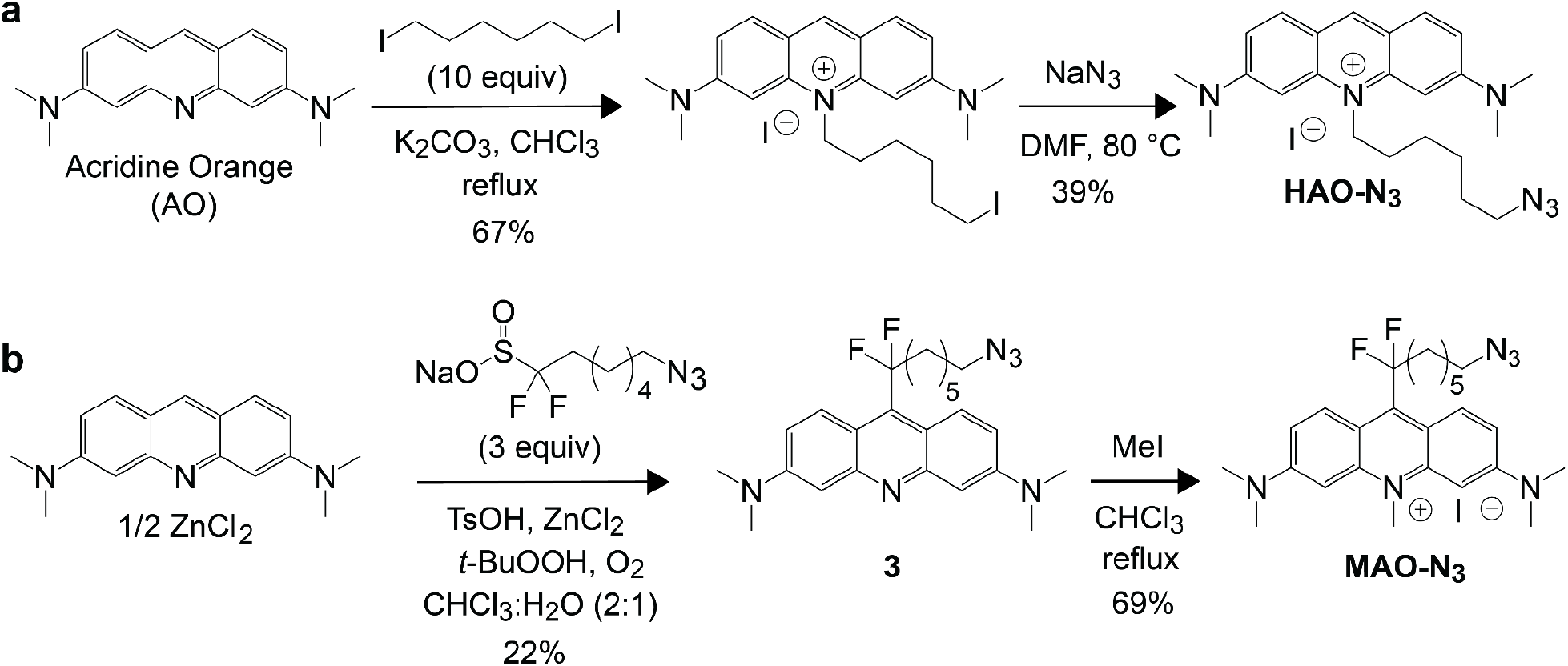
Scheme illustrating the synthetic steps used to prepare HAO-N_3_ and MAO-N_3_. Synthetic details are available in **Supplementary Information.**

**Extended Data Fig. 2.**
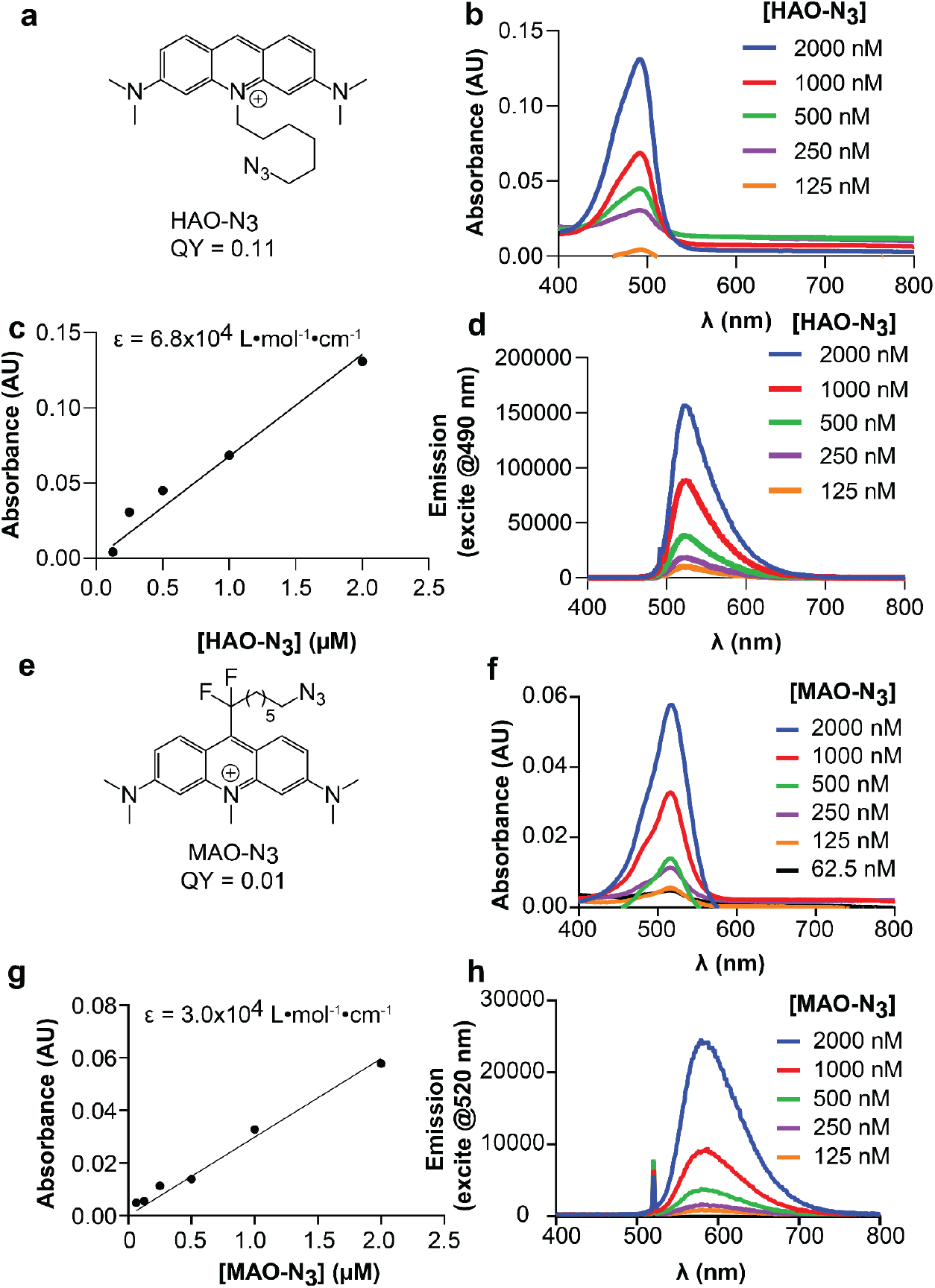
Photophysical properties of MAO-N_3_ and HAO-N_3_. **a,** Structure and quantum yield of HAO-N_3_. **b,** Concentration-dependent UV-Vis absorption curve of HAO-N_3_ shows a λ_max_ of 496 nm. **c,** Plot of absorbance @ λ_max_ against concentration of HAO-N_3_ and the calculated extinction coefficient. **d,** Concentration dependent fluorescence emission of HAO-N_3_ reveals a λ_em_ of 517 nm. **e,** Structure and quantum yield of MAO-N_3_. **f,** Concentration-dependent UV-Vis absorption curve of MAO-N_3_ shows a λmax of 520 nm. **g,** Plot of absorbance @ λmax against concentration of MAO-N_3_ and its extinction coefficient. **h,** Concentration dependent fluorescence emission of MAO-N_3_ reveals a λem of 570 nm.

**Extended Data Fig. 3.**
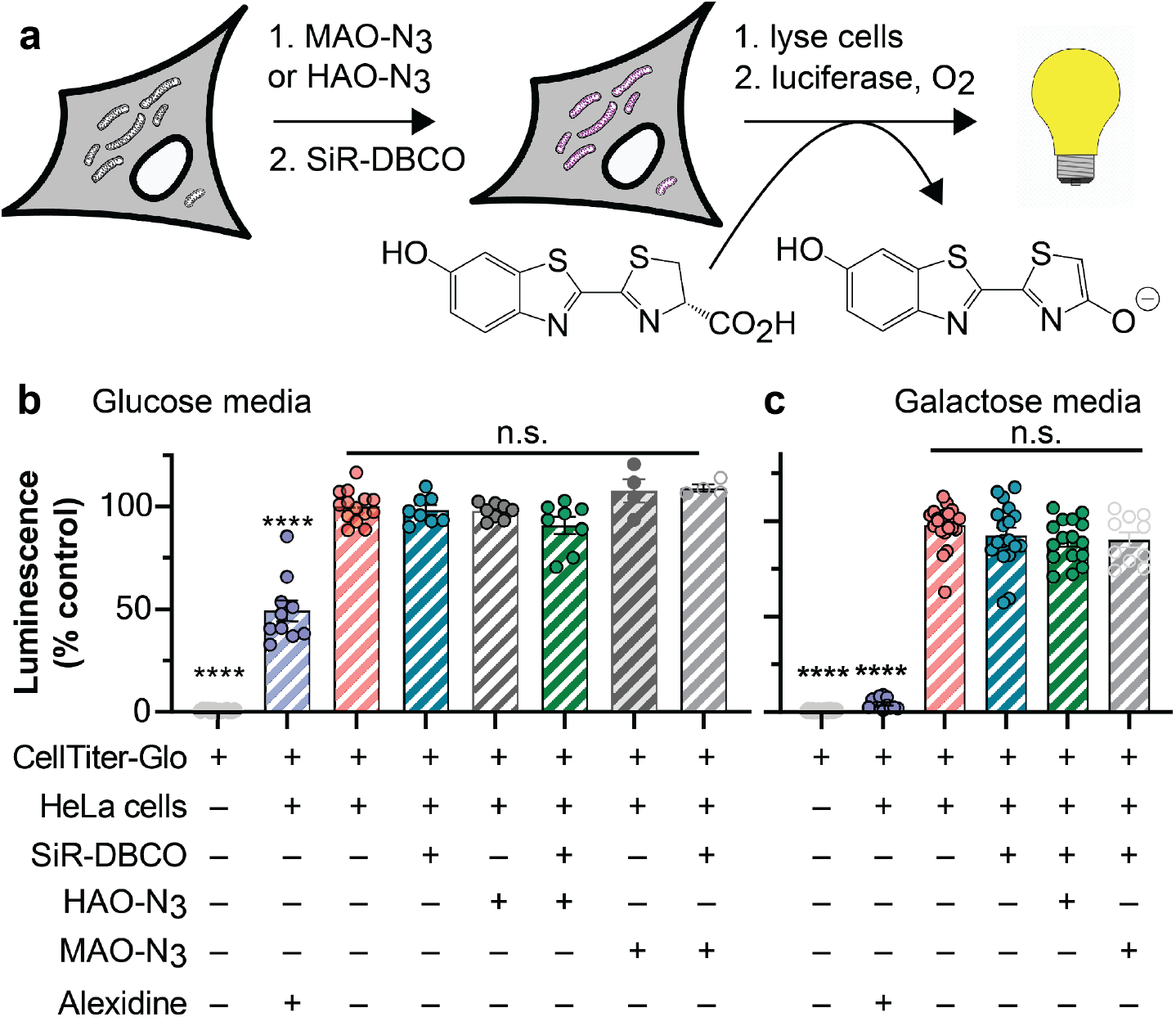
The HIDE probes HAO-SiR and MAO-SiR generated using HAO-N_3_ and MAO-N_3_ are non-toxic, even in galactose supplemented media. **a,** HeLa cells incubated in standard glucose (DMEM with 4.5 g/L glucose, supplemented with 10% FBS) or oxidative galactose-rich media (DMEM with 4.5 g/L galactose, supplemented with 10% FBS) were treated with HAO-N_3_ or MAO-N_3_ and SiR-DBCO and the ATP levels measured immediately as described in **Online Methods**. Plots show the relative bioluminescence signals (% relative to untreated cells). **b&c,** Plots show the relative bioluminescence signals of HeLa cell lysates incubated in **b,** standard or **c,** oxidative media. Untreated cells serve as a negative control; cells treated with 5 μM alexidine serve as a positive control. Alexidine inhibits the PTEN-like mitochondrial phosphatase PTPMT1 and disrupts mitochondrial integrity^29^. Error bars represent the standard error of measurement (sem); (****) p < 0.0001, from one-way ANOVA with Dunnett’s post-analysis accounting comparison to the negative control where the cells were incubated with cell culture media only (n.s. = not significant). Each bar represents the average of 12 biological replicates.

**Extended Data Fig. 4.**
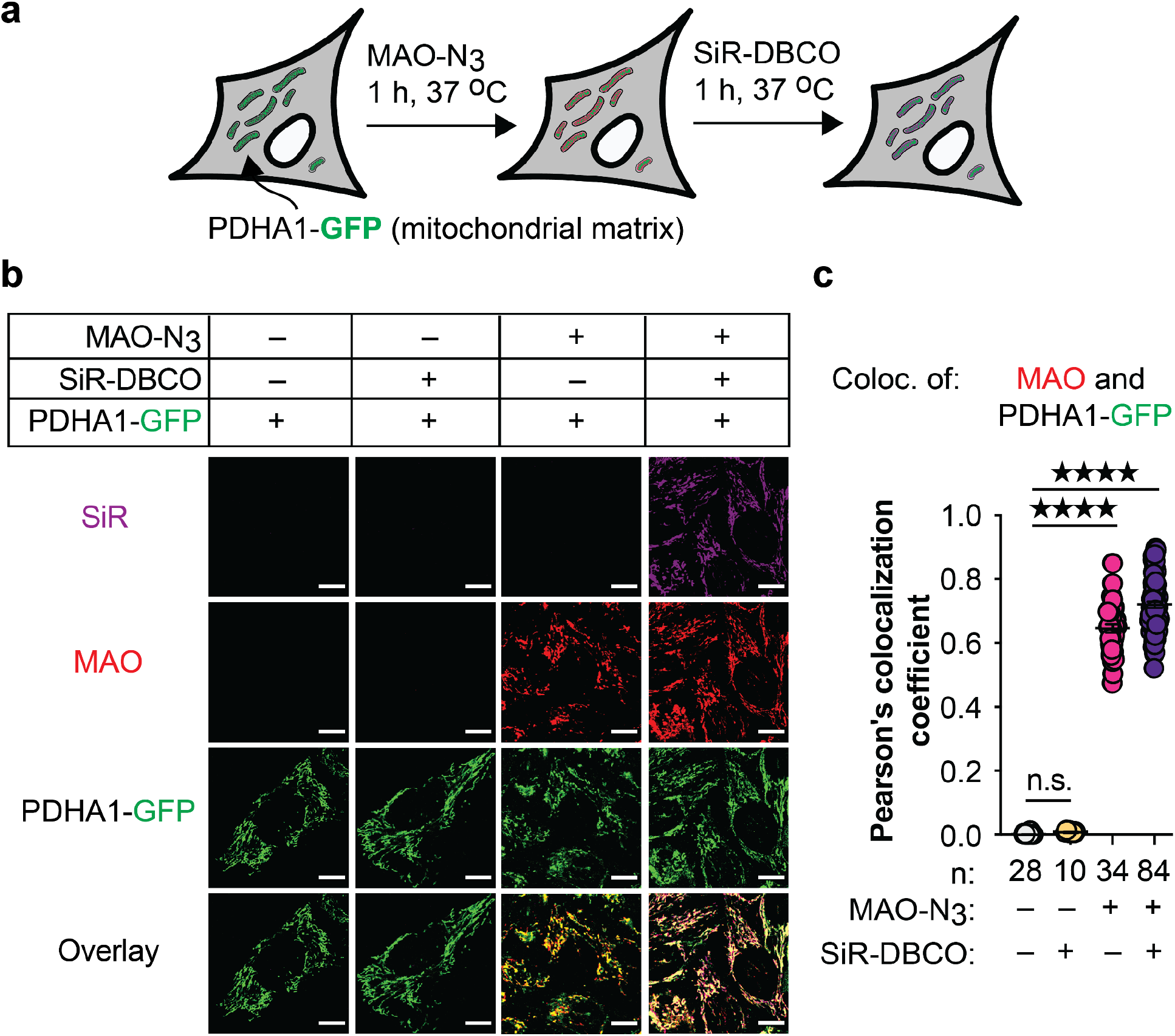
MAO-SiR, the HIDE probe generated from MAO-N_3_ and SiR-DBCO, localizes to mitochondria. **a,** HeLa cells expressing the mitochondrial matrix marker PDHA1-GFP (CellLight™ Mitochondria-GFP, BacMam 2.0) were treated as described in **Online Methods**. Cells were visualized using a point-scanning confocal microscope. **b,** Confocal images representing the colocalization of signal from MAO or SiR with PDHA1-GFP. **c,** Pearson’s colocalization coefficients PCC (MAO/PDHA1-GFP) = 0.77 ± 0.01. *n* represents the number of cells analyzed from at least two biological replicates. Scale bar: 10 μm. Error bars = s.e.m. **** p<0.0001, *** p<0.001, ** p<0.01, *p<0.1, n.s. not significant, from one-way ANOVA with Tukey’s multiple comparison test.

**Extended Data Fig. 5.**
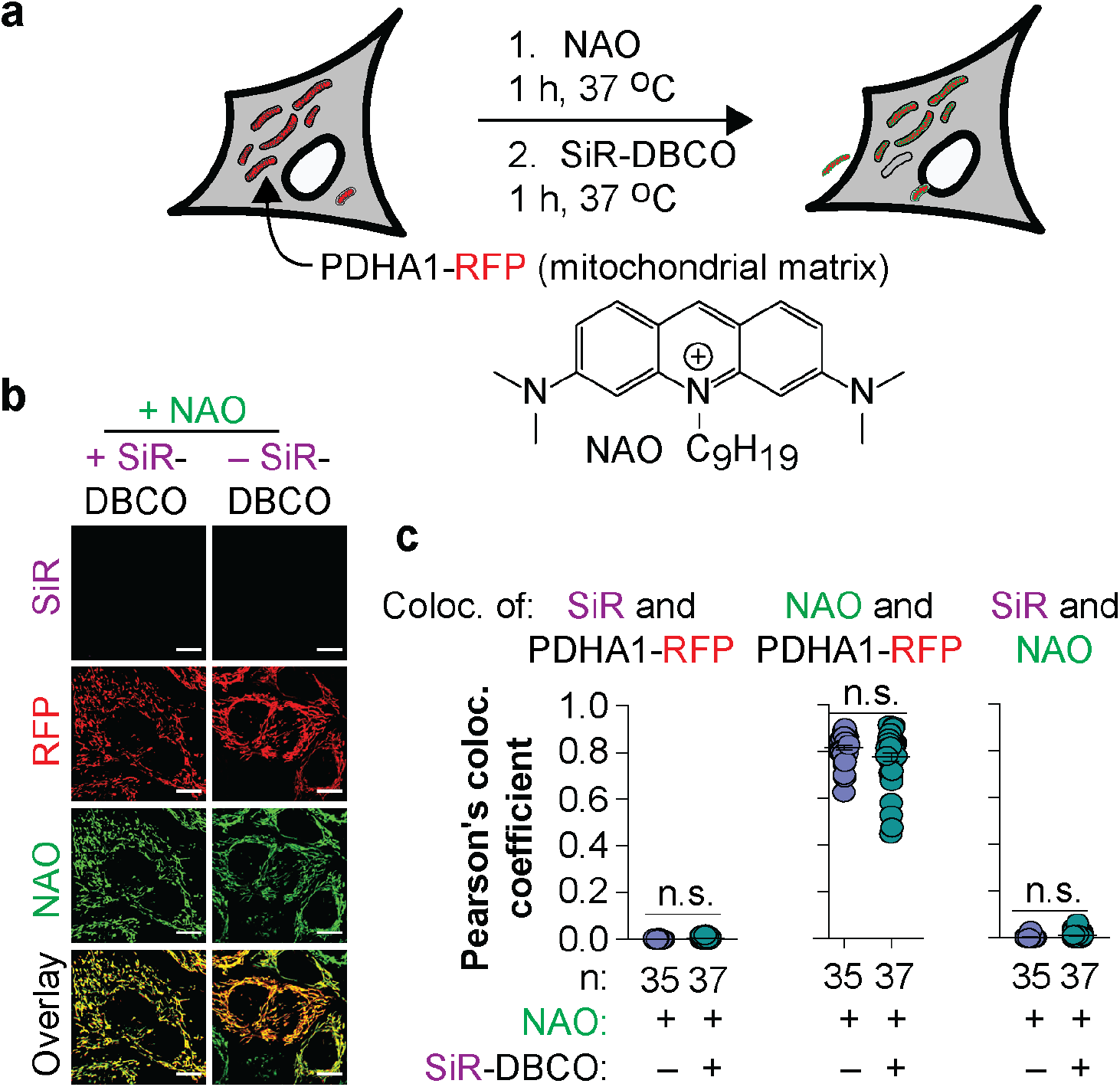
The azido group of MAO-N_3_ and HAO-N_3_ recruits SiR-DBCO to the mitochondria. **a,** HeLa cells expressing the mitochondrial matrix marker PDHA1-RFP (CellLight™ Mitochondria-RFP, BacMam 2.0) were treated with NAO and SiR-DBCO as described in **Online Methods** and visualized using a point-scanning confocal microscope. **b,** Representative confocal images used for quantification of Pearson’s colocalization coefficients. **c,** Pearson’s colocalization coefficients (PCC) representing the colocalization of signal from NAO or SiR with mitochondria marker PDHA1-RFP in HeLa cells. PCC (NAO/PDHA1-RFP) = 0.81 ± 0.01; no significant signal was observed in the SiR channel. Scale bar: 10 μm. n = # of cells. Error bars = s.e.m. ****p<0.0001, ***p<0.001, **p<0.01, *p<0.1, n.s. not significant, from one-way ANOVA with Tukey’s multiple comparison test.

**Extended Data Fig. 6.**
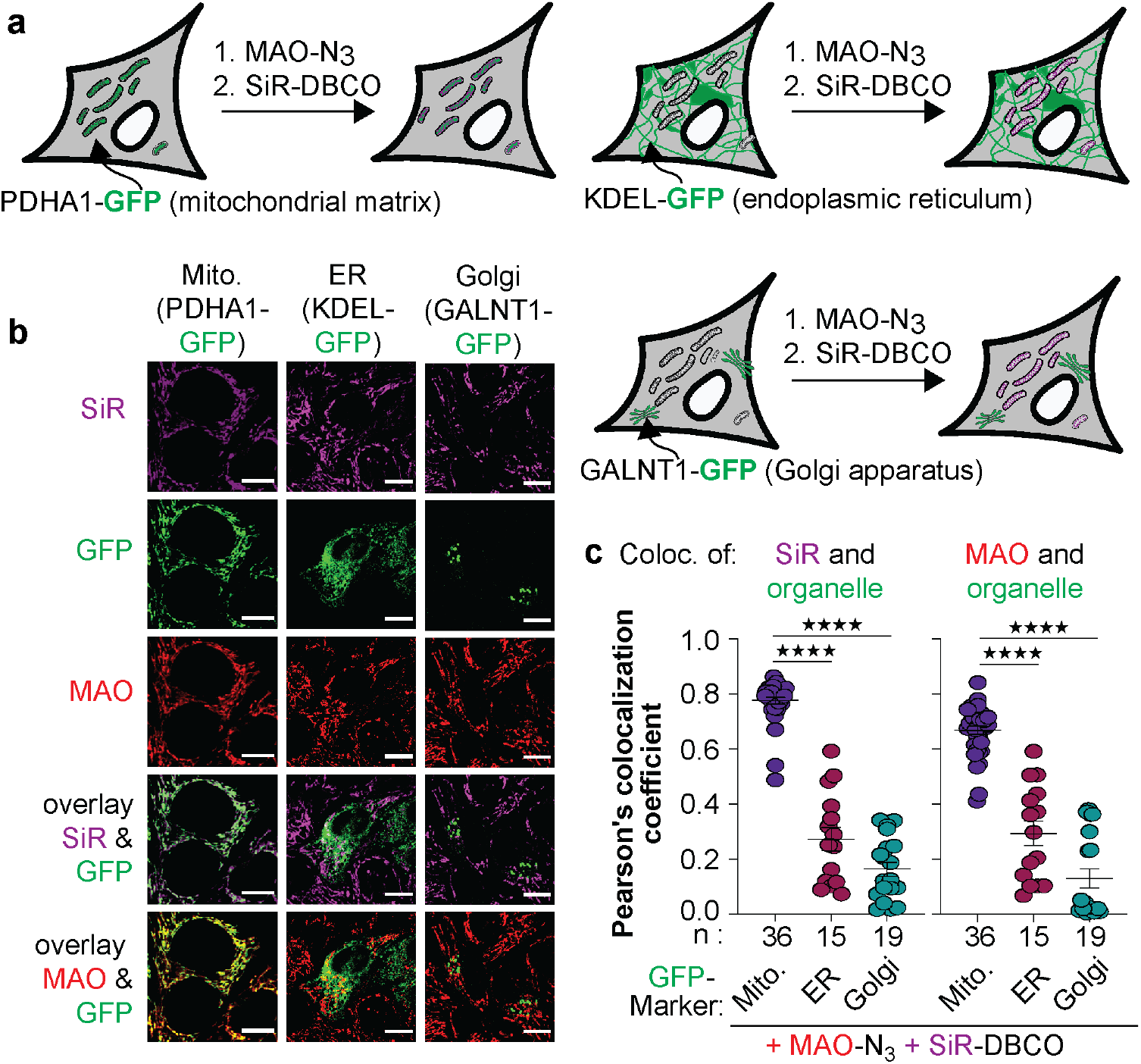
MAO-SiR, the HIDE probe generated from MAO-N_3_ and SiR-DBCO, does not localize significantly to the ER or Golgi. **a,** HeLa cells expressing the mitochondrial matrix marker PDHA1-GFP (CellLight™ Mitochondria-GFP, BacMam 2.0), the endoplasmic reticulum (ER) marker KDEL-GFP (CellLight™ ER-GFP, BacMam 2.0), or the Golgi marker GALNT1-GFP (CellLight™ Golgi-GFP, BacMam 2.0) were treated as described in **Online Methods** and imaged on a point-scanning confocal microscope. **b,** Representative images and **c,** Pearson’s colocalization coefficients (PCC) quantifying the colocalization between signals from MAO and SiR with PDHA1-GFP, KDEL-GFP, and GALNT1-GFP, respectively. PCC (MAO/PDHA1-GFP) = 0.66 ± 0.01, PCC(SiR/PDHA1-GFP) = 0.66 ± 0.01; PCC (MAO/GALNT1-GFP) = 0.16 ± 0.03; PCC (SiR/GALNT1-GFP) = 0.13 ± 0.03; PCC (MAO/KDEL-GFP) = 0.27 ± 0.04; PCC (SiR/KDEL-GFP) = 0.29 ± 0.04. Scale bar: 10 μm. *n* represents of cells. Error bars = s.e.m. ****p<0.0001, ***p<0.001, **p<0.01, *p<0.1, n.s. not significant, from one-way ANOVA with Tukey’s multiple comparison test.

**Extended Data Fig. 7.**
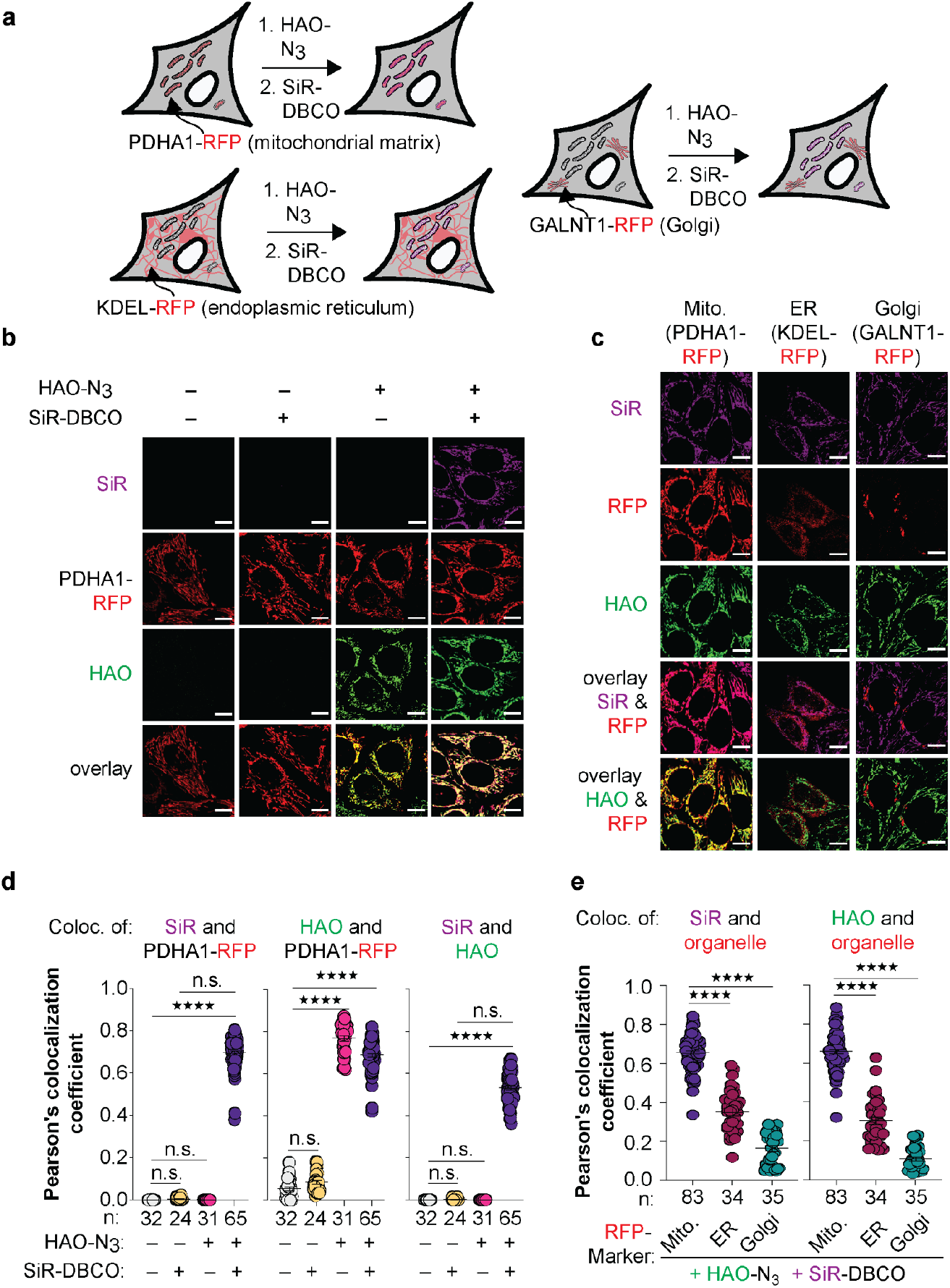
HAO-SiR, the HIDE probe generated from HAO-N_3_ and SiR-DBCO, localizes to mitochondria. **a,** HeLa cells expressing the mitochondrial matrix marker PDHA1-RFP (CellLight™Mitochondria-RFP, BacMam 2.0), the endoplasmic reticulum (ER) marker KDEL-RFP (CellLight™ ER-RFP, BacMam 2.0), or the Golgi marker GALNT1-RFP (CellLight™ Golgi-RFP, BacMam 2.0) were treated as described in **Online Methods** and imaged on a point-scanning confocal microscope. **b-c,** Confocal images representing the colocalization signal from HAO-N_3_ or SiR-DBCO with **b**, mitochondria marker PDHA1-RFP or **c**, ER marker KDEL-RFP and Golgi marker GALNT1-RFP in HeLa cells. **d-e,** Pearson’s colocalization coefficients (PCC) quantifying the colocalization of signal between HAO or SiR and mitochondria, ER or Golgi markers in HeLa cells. PCC(SiR/PDHA1-RFP) = 0.70 ± 0.01, PCC (HAO/PDHA1-RFP) = 0.69 ± 0.01, PCC (SiR/HAO) = 0.53 ± 0.01; PCC (HAO/GALNT1-RFP) = 0.11 ± 0.01; PCC (SiR/GALNT1-RFP) = 0.17 ± 0.01; PCC(HAO/KDEL-RFP) = 0.31 ± 0.02; PCC(SiR/KDEL-RFP) = 0.35 ± 0.02. Scale bar: 10 μm. n = # of cells. Error bars = s.e.m. ****p<0.0001, ***p<0.001, **p<0.01, *p<0.1, n.s. not significant, from one-way ANOVA with Tukey’s multiple comparison test.

**Extended Data Fig. 8.**
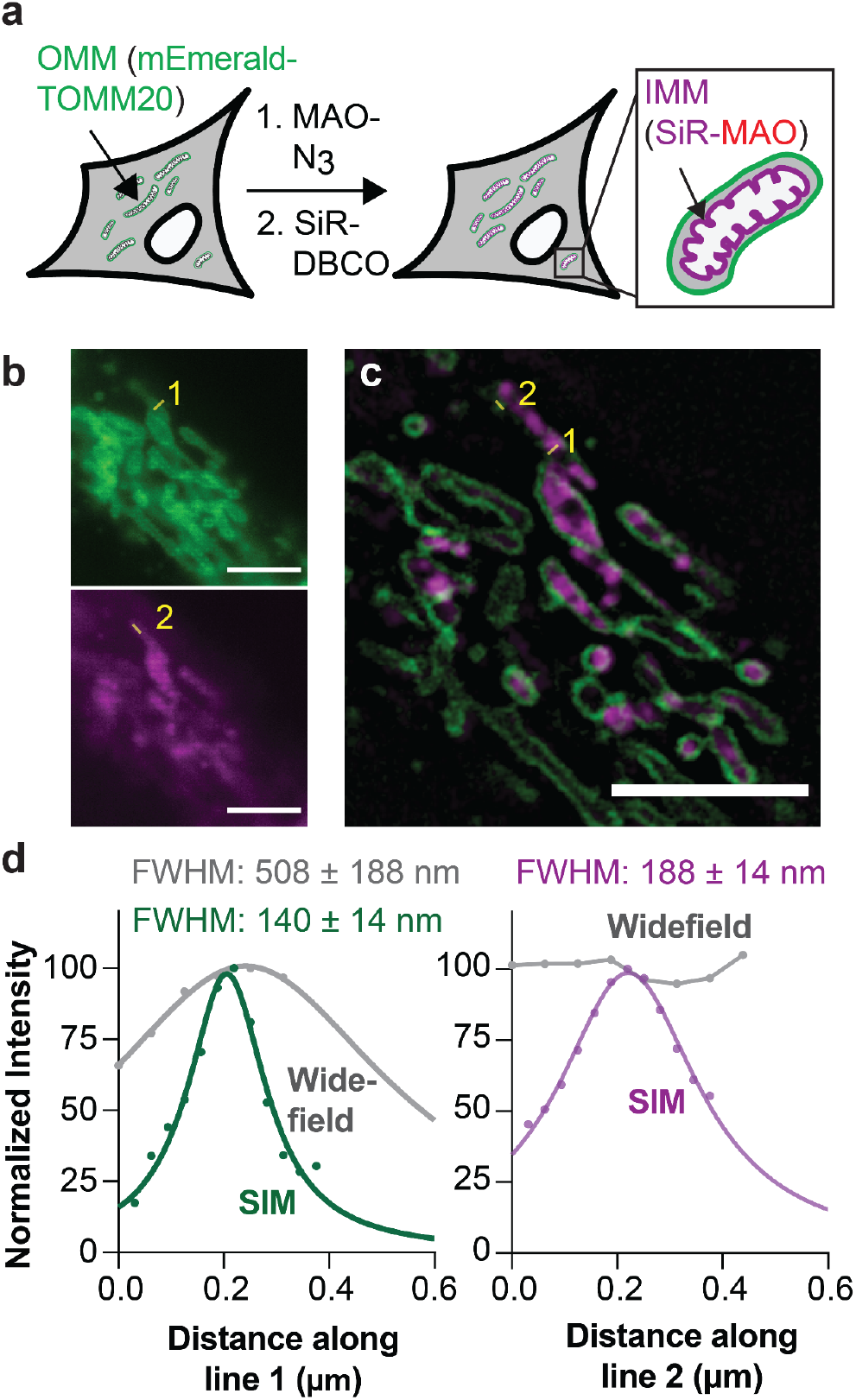
Two-color SIM imaging of mitochondria using the HIDE probe MAO-SiR and the OMM marker mEmerald-TOMM20. **a,** HeLa cells expressing mEmerald-TOMM20 were treated with MAO-N_3_ and SiR-DBCO as described in **Online Methods** and imaged using lattice SIM. **b,** Widefield images of HeLa cells treated as described in **a**; mEmerald-TOMM20 (TOP, green, 510 nm) and SiR (BOTTOM, magenta, 660 nm) channels. **c,** The reconstructed SIM image showing the merger of SiR and mEmerald signals. **d,** Plot illustrating the fit of signals along line **1** (LEFT) in panels **b** and **c** to a Gaussian function to establish FWHM values of 508 ± 188 nm for the widefield image and 140 ± 14 nm for the SIM image. Plot illustrating the fit of signals along line 2 (RIGHT) in panels **b** and **c** to a Gaussian function to establish a FWHM of 188 ± 14 nm for the SIM image.

**Extended Data Fig. 9.**
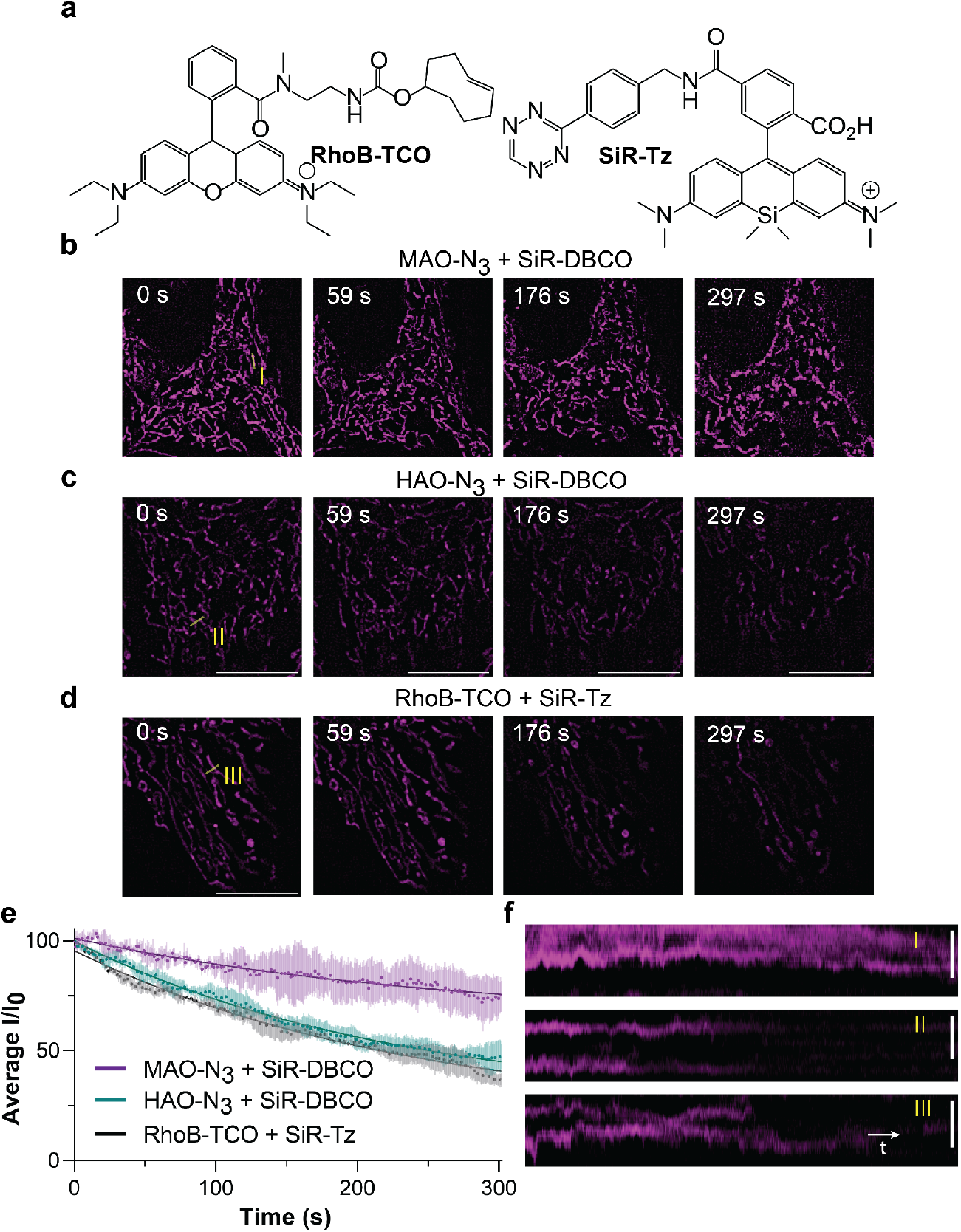
The IMM-selective HIDE probe supports extended-time SIM imaging in live cells. **a,** Structures of RhoB-TCO and SiR-Tz. HeLa cells were treated with **b,** MAO-N_3_, **c,** HAO-N_3_ or **d,** RhoB-TCO probes followed by SiR-DBCO (**b&c**) or SiR-Tz (**d**) as described in **Online Methods,** and imaged on a widefield microscope equipped with lattice SIM. Scale bar: 10 μm. **e,** Plot illustrating normalized SiR fluorescence signals of MAO-SiR, HAO-SiR and RhoB-SiR over time. t_1/2_ (HAO-SiR) = 248.2 ± 1.2 s; t_1/2_ (RhoB-SiR) = 233.0 ± 1.3 s. *n* = 3 regions of interest (ROI), *N* = 1 cell. **f,** kymograph of line I (MAO-SiR), II (HAO-SiR), and III (RhoB-SiR). Scale bar: 1 μm.

**Extended Data Fig. 10.**
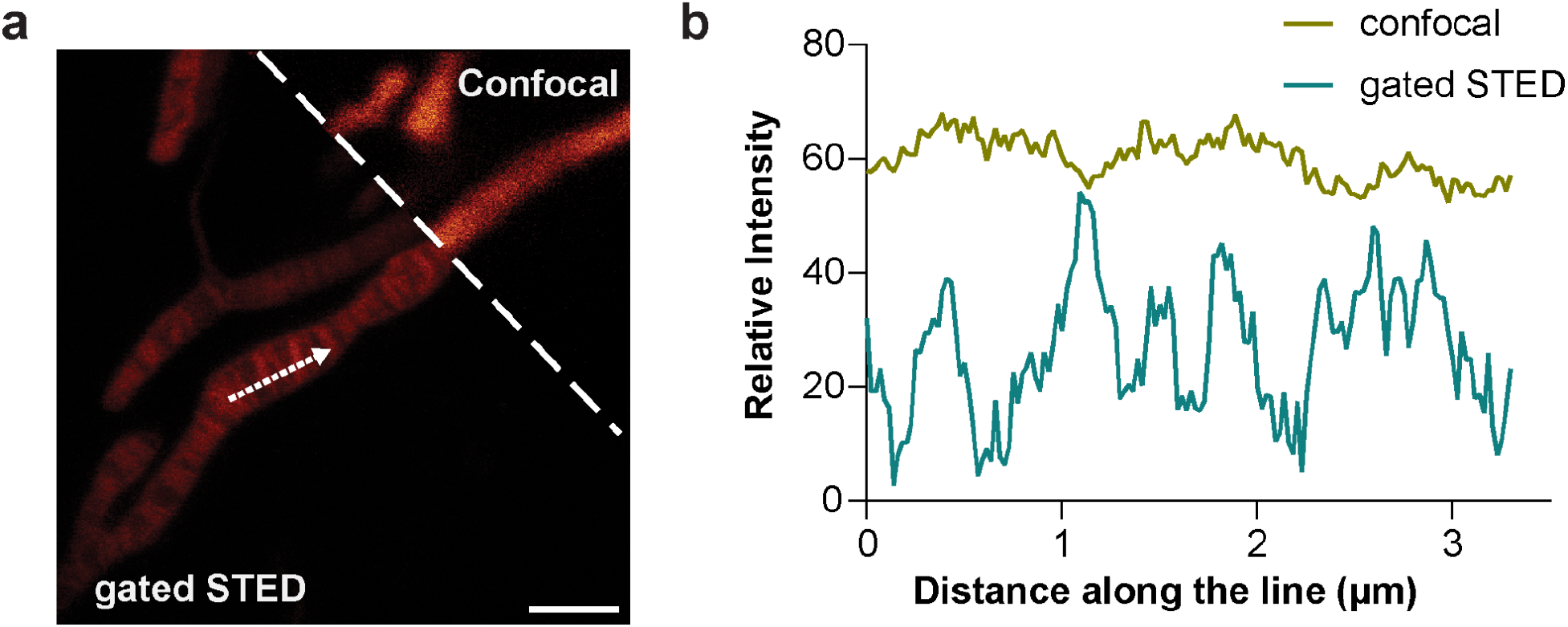
Comparison of diffraction limited confocal microscopy vs. time-gated STED microscopy when imaging the IMM using MAO-SiR. **a,** gated STED and confocal images of signals due to MAO-SiR. Scale bar: 2 μm. **b,** Plots of signals due to SiR in confocal and STED channels along the dotted arrow in **a**.

