## Supplementary Information for "Long-term, super-resolution HIDE imaging of the inner mitochondrial membrane in live cells with a cell-permeant lipid probe"

**Affiliations:**

**Table of Contents**

I. Supplementary Figure

II. Synthesis of HAO-$N_3$, MAO-$N_3$ and $HMDS_{655}$-DBCO

   A. General Considerations

   B. Synthesis of HAO-$N_3$

   C. Synthesis of MAO-$N_3$

   D. Synthesis of $HMDS_{655}$-DBCO

III. NMR Spectra

IV. References

### I. Supplementary Figure

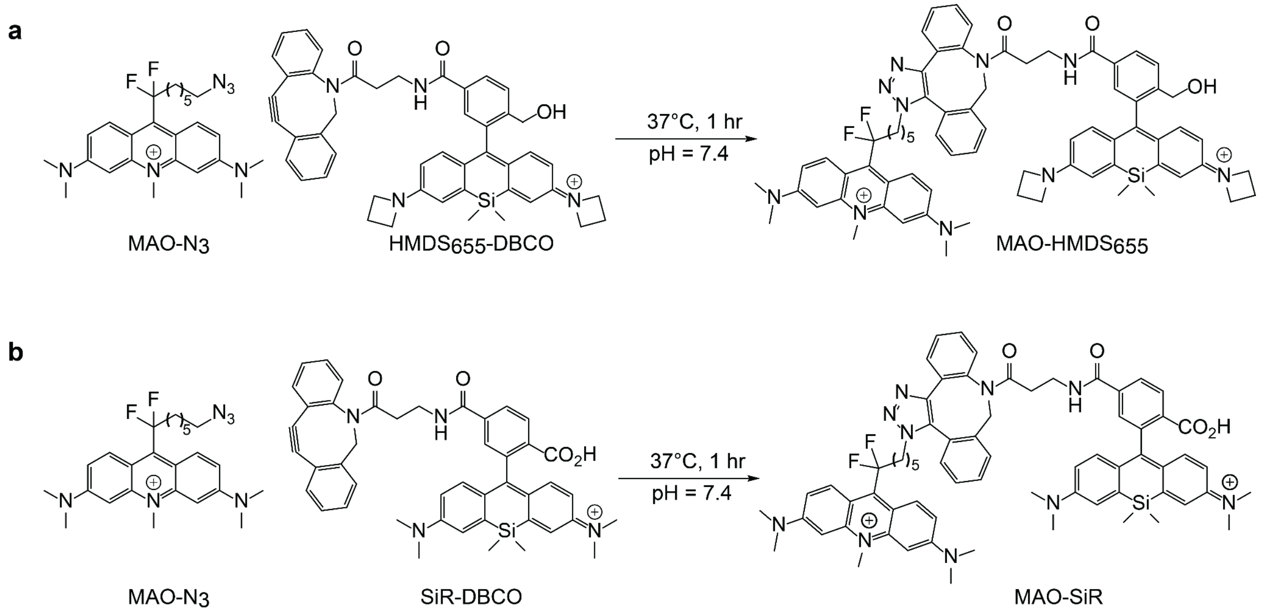


**Supplementary Fig 1. Inner mitochondrial membrane-specific HIDE probes MAO-HMDS$_{655}$ and MAO-SiR assembled using SPAAC reaction.**

### II. Synthesis of HAO-N$_3$, MAO-N$_3$ and HMDS$_{655}$-DBCO

#### A. General considerations

Unless otherwise noticed, all reactions were performed in a nitrogen atmosphere. Chemicals used for synthesis were either prepared according to literature reports or purchased from commercial sources and used without further purification. Flash chromatography was performed using a Teledyne Isco CombiFlash Rf system using pre-packed columns with RediSep Rf silica (40–60 μm). $^{1}$H NMR and $^{13}$C NMR spectra were recorded on Bruker Neo-500 (500 MHz for $^{1}$H, 126 MHz for $^{13}$C and 471 MHz for $^{19}$F) or AV-600 (600 MHz for $^{1}$H and 151 MHz for $^{13}$C) NMR spectrometers. The values of chemical shifts (δ) are reported in ppm relative to the solvent residual signals of $CD_3OD$ (3.31 ppm for $^{1}$H, 49.0 ppm for $^{13}$C), $CDCl_3$ (7.26 ppm for $^{1}$H, 77.2 ppm for $^{13}$C) or $CD_3CN$ (1.94 ppm for $^{1}$H, 1.32, 118.3 ppm for $^{13}$C). Coupling constants (J) are reported in Hz. High-resolution mass spectra (HRMS) were recorded on an Angilent QTOF LCMS with ESI connected to an Agilent UHPLC. HPLC purifications were performed on a reverse-phase column (YMC-Triart C18, 150 mm × 10 mm S-5 μm, 12 nm) using an Angilent HPLC system.

#### B. Synthesis of HAO-$N_3$

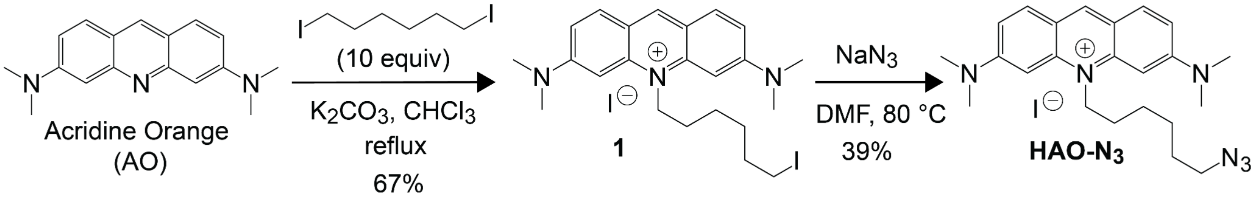


Acridine orange "free base" was obtained according to literature[1]. Briefly, 4 g acridine orange ½ $ZnCl_2$ (Chem-Impex) was dissolved 100 mL $H_2O$ and then treated with excess NaOH solution (4 N) for 30 min. The free acridine orange was extracted with $CH_2Cl_2$, washed with water, and dried over sodium sulfate. Removal of solvent under reduced pressure and drying under vacuum afforded 1.85 g of free acridine orange as a dark red-brown powder, which was used in the next step without further purification.

**Preparation of 3,6-bis(dimethylamino)-10-(6-iodohexyl)acridin-10-ium (1).** To a 20 mL microwave vial charged with $K_2CO_3$ (17.25 mg, 0.125 mmol, 0.5 equiv) was added acridine orange solution (92.5 mg, 0.25 mmol, 1 equiv in 5 mL $CHCl_3$), followed by addition of 1,6-diiodohexane (0.41 mL, 2.5 mmol, 10 equiv). The reaction mixture was heated at reflux for 16h, cooled, filtered, and concentrated *in vacuo*. The product **1** was obtained as a dark-red solid (100 mg, 67%), upon purification by flash column chromatography (neutral $Al_2O_3$, 0 to 5% MeOH:$CH_2Cl_2$). $^1$H NMR (500 MHz, $CDCl_3$) δ 8.68 (s, 1H), 7.89 (d, $J$ = 9.3 Hz, 2H), 7.05 (dd, $J$ = 9.3, 2.1 Hz, 2H), 6.67 (d, $J$ = 2.2 Hz, 2H), 4.99 – 4.89 (m, 2H), 3.25 (t, $J$ = 6.8 Hz, 2H), 2.01 (p, $J$ = 7.8 Hz, 2H), 1.93 – 1.82 (m, 2H), 1.80 – 1.70 (m, 2H), 1.63 – 1.58 (m, 2H). $^{13}$C NMR (126 MHz, $CDCl_3$) δ 155.7, 143.1, 142.8, 133.4, 117.3, 114.2, 93.2, 48.5, 41.5, 33.1, 30.5, 26.3, 26.2, 7.9. HRMS (ES+) calcd for ($C_{23}H_{31}IN_3$) [$M^+$] 476.1557, found 476.1549.

**Preparation of HAO-$N_3$.** Compound **1** (30.2 mg, 0.05 mmol, 1.0 equiv) and $NaN_3$ (9.8 mg, 3 equiv) were dissolved in DMF (1 mL) in an 8 mL microwave vial. The reaction was heated at 80 °C for 16 h and monitored by TLC. Upon completion, the reaction mixture was diluted with 30 mL $CH_2Cl_2$ and washed 3X with brine (30 mL), dried ($Na_2SO_4$) and evaporated. The corresponding product HAO-$N_3$ was obtained as a dark-red solid (10.2 mg, 39%), upon purification by flash column chromatography (neutral $Al_2O_3$, 0 to 10% MeOH:$CH_2Cl_2$). $^1$H NMR (500 MHz, Chloroform-*d*) δ 8.64 (s, 1H), 7.90 (d, $J$ = 9.3 Hz, 2H), 7.10 (dd, $J$ = 9.2, 2.2 Hz, 2H), 6.78 (d, $J$ = 2.3 Hz, 2H), 5.10 – 4.94 (m, 2H), 3.36 (s, 12H), 3.31 (t, $J$ = 6.7 Hz, 2H), 2.03 (tt, $J$ = 7.9 Hz, 2H), 1.76 (tt, $J$ = 7.6 Hz, 2H), 1.71 – 1.64 (m, 2H), 1.59 – 1.52 (m, 1H). $^{13}$C NMR (126

MHz, $CDCl_3$) δ 155.8, 143.3, 142.8, 133.5, 117.3, 114.2, 93.0, 51.5, 47.9, 41.1, 29.7, 28.9, 26.8, 26.3. HRMS (ES+) calcd for ($C_{23}H_{31}N_6$) [$M^+$] 391.2605, found 391.2594.

##### C. Synthesis of MAO-$N_3$

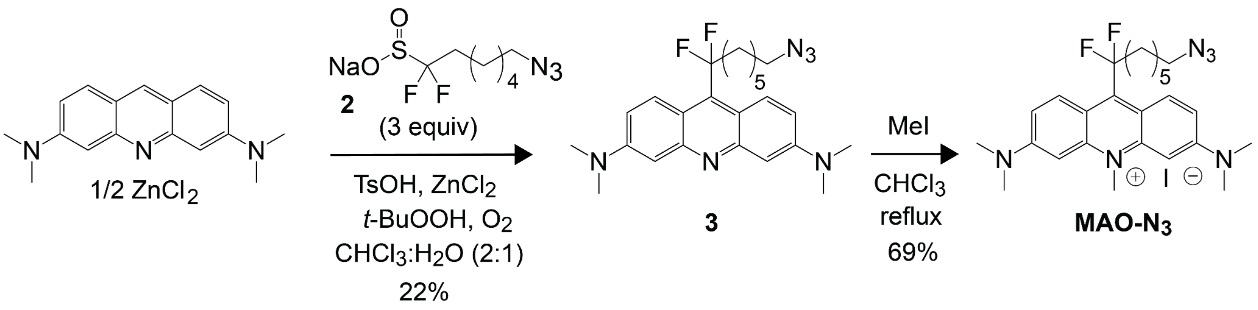


**Preparation of 9-(7-azido-1,1-difluoroheptyl)-*N*³,*N*³,*N*⁶,*N*⁶-tetramethylacridine-3,6-diamine (3).** Azide **3** was synthesized as reported[2]. Difluoride **2** (90.0 mg, 0.34 mmol, 4.0 equiv)[2], acridine orange hemi $ZnCl_2$ salt (34.8 mg, 0.09 mmol, 1.0 equiv ) and $ZnCl_2$ (17.5 mg, 0.13 mmol, 1.5 equiv) was dissolved in $CHCl_3$ (1 mL) and $H_2O$ (0.5 mL) in a 4 mL vial equipped with septa screw cap, followed by addition of TsOH•$H_2O$ (32.5 mg, 0.17 mmol, 2 equiv). The reaction mixture was cooled to 0 °C and *tert*-butyl hydroperoxide (TBHP, 70% in water, 0.07 ml, 60.5 mg, 0.47 mmol, 5.5 equiv) was added dropwise with vigorous stirring over 5 min. The reaction was then heated at 50 °C for 1 h, cooled to RT, and stirred for 24 h under continuous air flow. The mixture was then diluted with $CH_2Cl_2$ (2 mL), separated, and the aqueous layer was washed with $CH_2Cl_2$ (2 x 2 mL). The combined organic layers were washed with $NaHCO_3$, brine, dried ($Na_2SO_4$), and concentrated *in vacuo*. Azide **3** was obtained as a deep-red sticky solid (10 mg, 22%), upon purification by flash column chromatography (neutral $Al_2O_3$, 0 to 5% MeOH:$CH_2Cl_2$). The NMR spectrum of the obtained product matched the literature report.

**Preparation of**
**9-(7-azido-1,1-difluoroheptyl)-3,6-bis(dimethylamino)-10-methylacridin-10-ium (MAO-$N_3$)**.
To an 8 mL microwave vial was added a $CHCl_3$ (3 mL) solution of azide **3** (10 mg, 0.02 mmol, 1.0 equiv) and then $CH_3I$ (32.2 mg, 0.23 mmol, 10 equiv). The reaction mixture was heated to reflux overnight before cooling and purification *via* flash column chromatography (neutral $Al_2O_3$, 0 to 5% MeOH:$CH_2Cl_2$). The corresponding product **MAO-$N_3$** was obtained as a deep red solid (9.1 mg, 69%). $^1H$ NMR (600 MHz, Chloroform-*d*) δ 8.20 (d, *J* = 9.9 Hz, 2H), 7.12 (dd, *J* = 10.0, 2.5 Hz, 2H), 6.89 (d, *J* = 2.5 Hz, 2H), 4.54 (s, 3H), 3.41 (s, 12H), 3.30 (t, *J* = 6.8 Hz, 2H), 2.52 (tt, *J* = 17.1, 8.2 Hz, 2H), 1.83 – 1.74 (m, 2H), 1.63 (tt, *J* = 7.1, 6.8 Hz, 2H), 1.53 – 1.42 (m, 4H).

$^{13}C$ NMR (151 MHz, Chloroform-*d*) δ 154.4, 145.3 (t, *J* = 24.3 Hz), 144.6, 129.6 (t, *J* = 10.2 Hz), 125.1 (t, *J* = 247.4 Hz), 115.0, 114.5, 94.7, 51.4, 41.4, 41.3, 40.6 (t, *J* = 24.6 Hz), 28.8, 28.7, 26.6, 22.1. $^{19}F$ NMR (376 MHz, Chloroform-*d*) δ -81.48 (t, *J* = 17.4 Hz). HRMS (ES+) calcd for ($C_{23}H_{33}F_2N_6$) [$M^+$] 455.2729, found 455.2719.

##### D. Synthesis of HMDS$_{655}$-DBCO

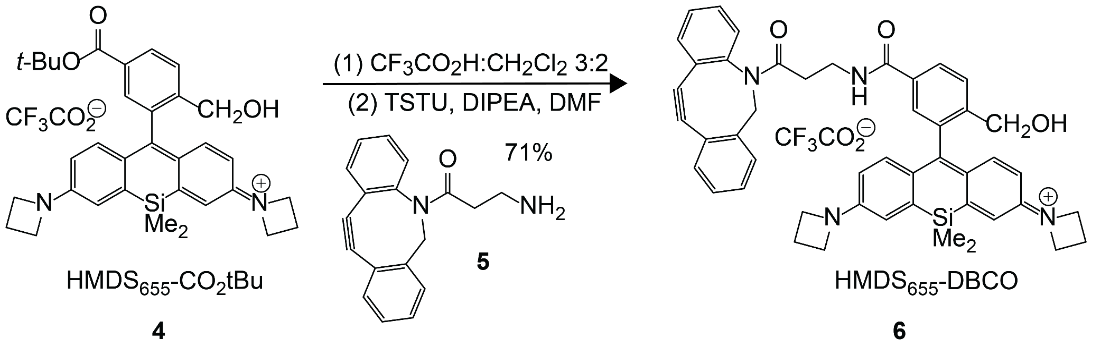


4 5 6

**Preparation of HMDS$_{655}$-CO$_2$tBu 4**. Compound **4** was synthesized according to a previous report[3].

**Preparation of HMDS$_{655}$-DBCO.** To a 4 mL vial equipped with a septa screw cap was added **5** (11.6 mg, 0.02 mmol, 1 equiv) and 2 mL of a 3:2 mixture of $CF_3CO_2H$ and $CH_2Cl_2$. The resulting reaction mixture was stirred for 2 h and monitored by TLC. Upon completion, the mixture was concentrated *in vacuo* and thoroughly dried under high vacuum. The product was redissolved in DMF (2 mL), DIPEA (13.9 mg, 0.11 mmol, 5.0 equiv) was added, and the reaction mixture stirred for 5 mins. A solution of TSTU (8.1 mg, 0.03 mmol, 1.25 equiv) and DIPEA (13.9 mg, 0.11 mmol, 5.0 equiv) in DMF (1 mL) was then added. The reaction was stirred overnight and purified using an Agilent semi-prep HPLC system on a reverse-phase column (YMC-Triart C18, 150 mm × 10 mm S-5 μm, 12 nm) with eluents A ($H_2O$ with 1% TFA) and B($CH_3CN$ with 1% TFA). The corresponding product HMDS$_{655}$-DBCO (**6**) was obtained as a blue solid (10.9 mg, 71%). $^1H$ NMR (500 MHz, MeOD) δ 7.64 (d, *J* = 7.8 Hz, 1H), 7.58 (s, 1H), 7.49 – 7.39 (m, 4H), 7.32 – 7.26 (m, 2H), 7.12 – 7.09 (m, 1H), 7.08 – 7.04 (m, 2H), 6.82 – 6.77 (m, 2H), 6.71 (dd, *J* = 8.1, 2.7 Hz, 2H), 6.44 (ddd, *J* = 8.6, 5.8, 2.6 Hz, 2H), 5.42 (s, 2H), 5.14 (d, *J* = 14.0 Hz, 2H), 3.92 – 3.79 (m, 8H), 3.69 (s, 2H), 2.48 (dq, *J* = 17.2, 5.9 Hz, 2H), 2.40 – 2.27 (m, 4H), 0.60 (s, 3H), 0.48 (s, 3H). $^{13}C$ NMR (151 MHz, MeOD) δ 173.48, 169.46, 152.59, 152.41, 152.30, 152.25, 149.33, 140.71, 140.61, 139.24, 135.19, 134.66, 133.34, 130.35, 129.92, 129.68, 129.62, 129.02, 128.82, 128.16, 127.77, 126.56, 124.53, 124.20, 123.65, 121.76, 119.23,

117.29, 116.49, 115.61, 115.34, 114.55, 114.50, 108.88, 94.23, 74.10, 56.51, 53.69, 53.66, 37.43, 35.36, 17.91, 0.15, -0.18.HRMS (ES+) calcd for ($C_{47}H_{45}N_4O_3Si^+$) [$M^+$] 741.3255, found 741.3261.

**III. NMR Spectra**

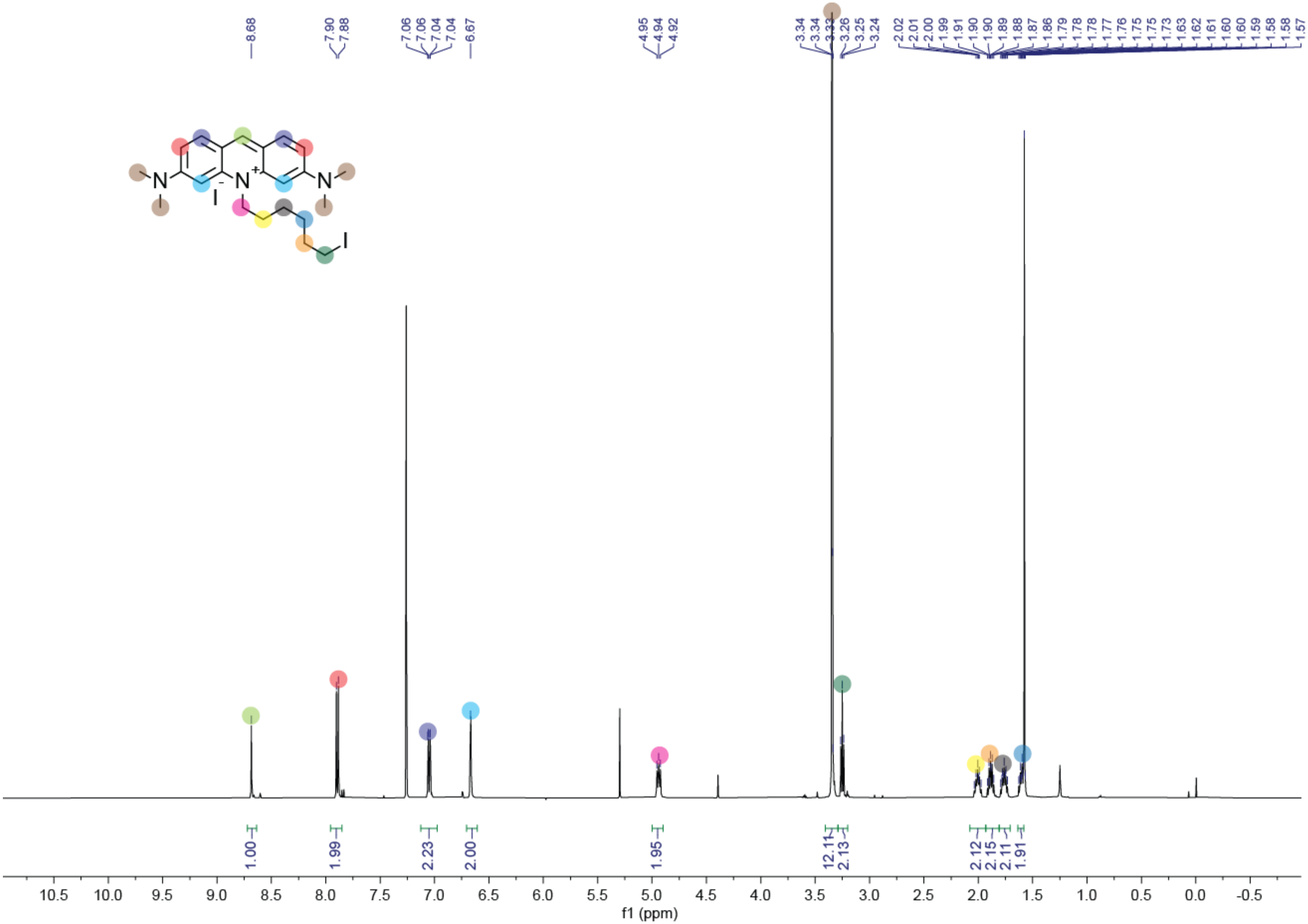
8.68
7.90
7.88
7.06
7.06
7.04
7.04
6.67
4.95
4.94
4.92
3.34
3.34
3.33
3.26
3.25
3.24
2.02
2.01
2.00
1.99
1.91
1.90
1.90
1.89
1.88
1.87
1.86
1.79
1.78
1.78
1.77
1.76
1.75
1.75
1.73
1.63
1.62
1.61
1.60
1.60
1.59
1.58
1.58
1.57
N
N
N
I
I
1.00
1.99
2.23
2.00
1.95
12.11
2.13
2.12
2.15
2.11
1.91
10.5
10.0
9.5
9.0
8.5
8.0
7.5
7.0
6.5
6.0
5.5
5.0
4.5
4.0
3.5
3.0
2.5
2.0
1.5
1.0
0.5
0.0
-0.5
f1 (ppm)

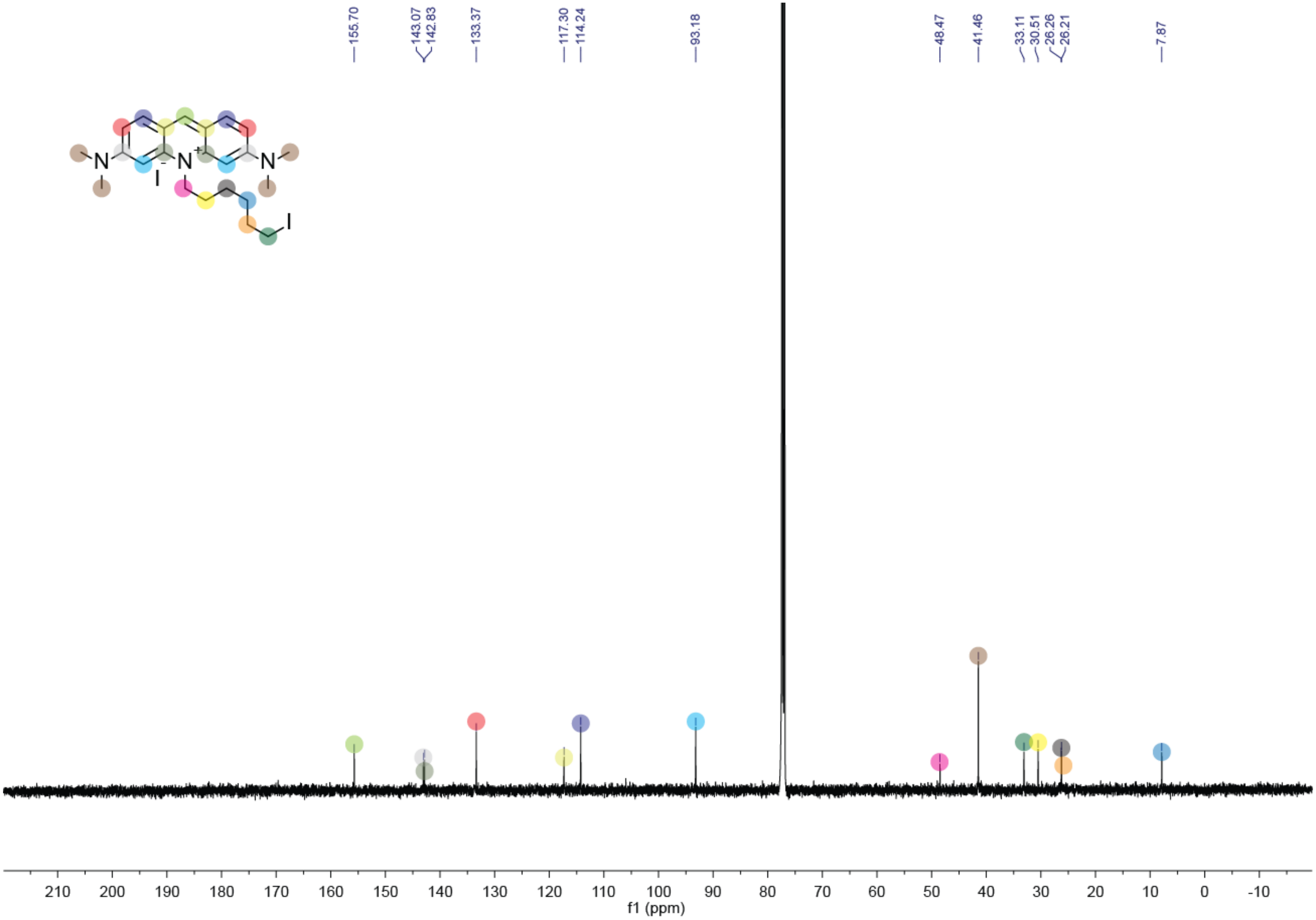

155.70
143.07
142.83
133.37
117.30
114.24
93.18
48.47
41.46
33.11
30.51
26.26
26.21
7.87
N
N
N
+
I
-
I
210 200 190 180 170 160 150 140 130 120 110 100 90 80 70 60 50 40 30 20 10 0 -10
f1 (ppm)

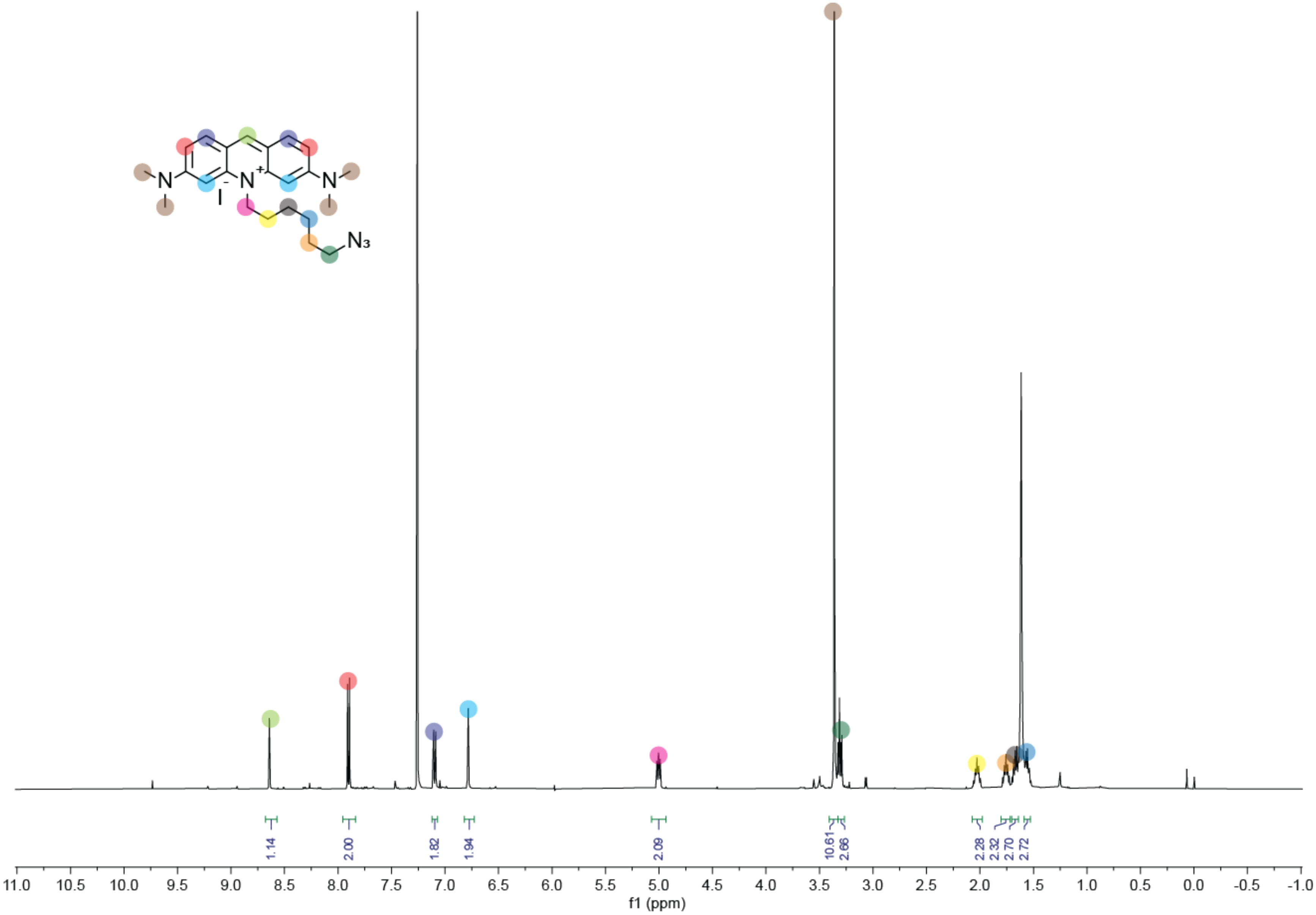
N
N
N
I
N3
1.14
2.00
1.82
1.94
2.09
10.61
2.66
2.28
2.32
2.70
2.72
11.0 10.5 10.0 9.5 9.0 8.5 8.0 7.5 7.0 6.5 6.0 5.5 5.0 4.5 4.0 3.5 3.0 2.5 2.0 1.5 1.0 0.5 0.0 -0.5 -1.0
f1 (ppm)

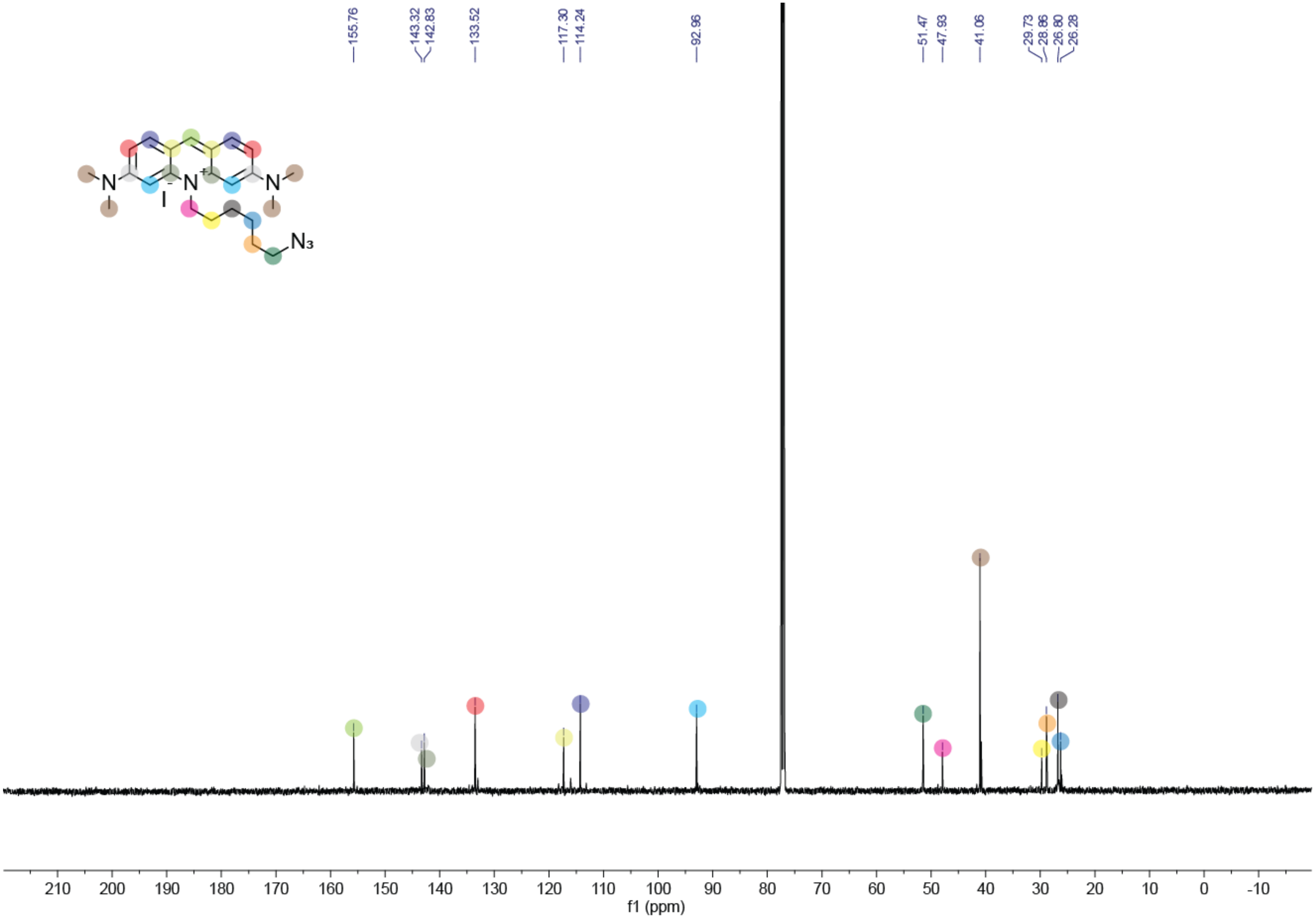

155.76
143.32
142.83
133.52
117.30
114.24
92.96
51.47
47.93
41.06
29.73
28.86
26.80
26.28
N
N+
N
I-
N3
210 200 190 180 170 160 150 140 130 120 110 100 90 80 70 60 50 40 30 20 10 0 -10
f1 (ppm)

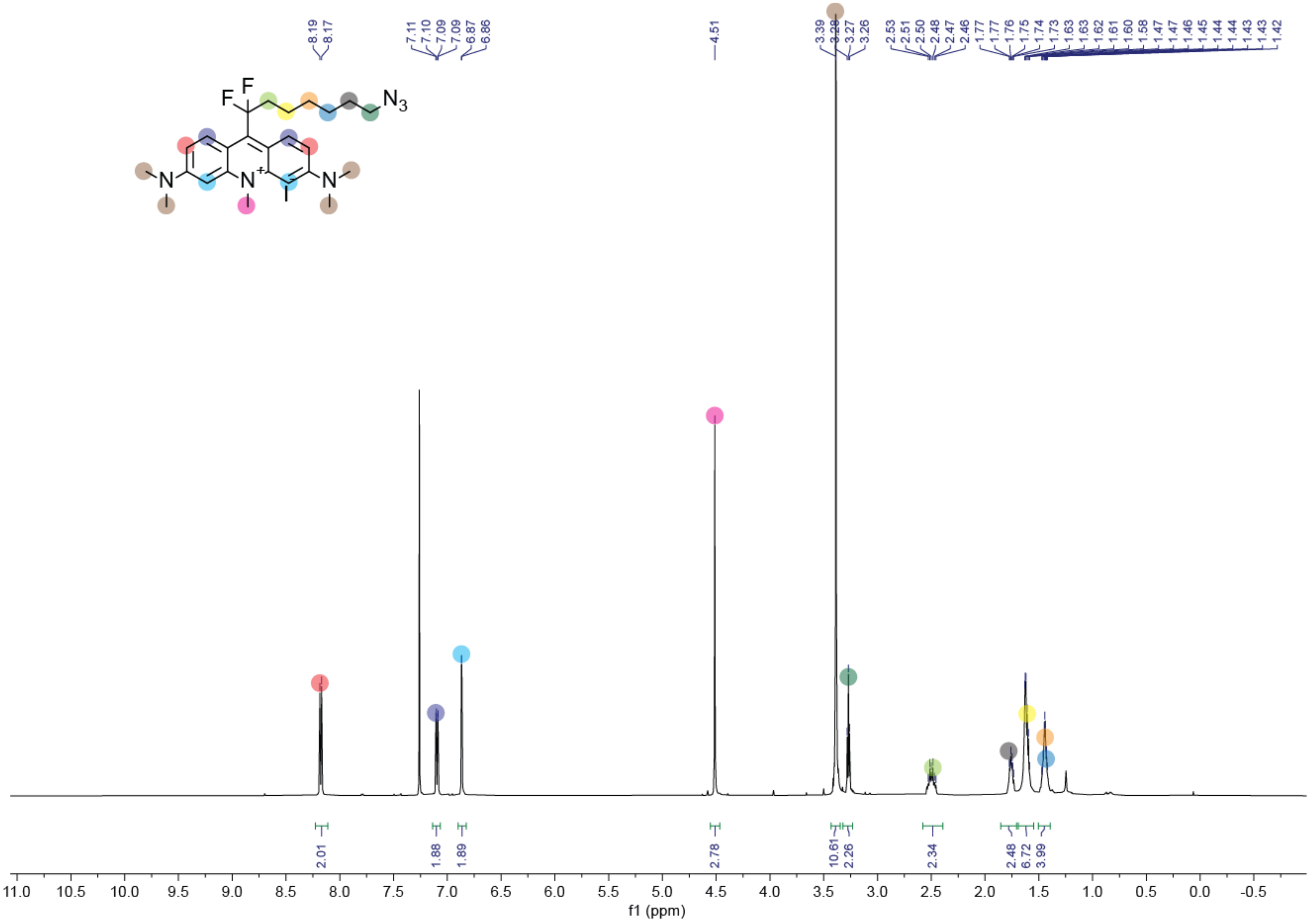

8.19
8.17
7.11
7.10
7.09
7.09
6.87
6.86
4.51
3.39
3.28
3.27
3.26
2.53
2.51
2.50
2.48
2.47
2.46
1.77
1.76
1.75
1.74
1.73
1.63
1.63
1.62
1.61
1.60
1.58
1.47
1.47
1.46
1.45
1.44
1.44
1.43
1.43
1.42
F
F
N3
N
N+
N
2.01
1.88
1.89
2.78
10.61
2.26
2.34
2.48
6.72
3.99
11.0
10.5
10.0
9.5
9.0
8.5
8.0
7.5
7.0
6.5
6.0
5.5
5.0
4.5
4.0
3.5
3.0
2.5
2.0
1.5
1.0
0.5
0.0
-0.5
f1 (ppm)

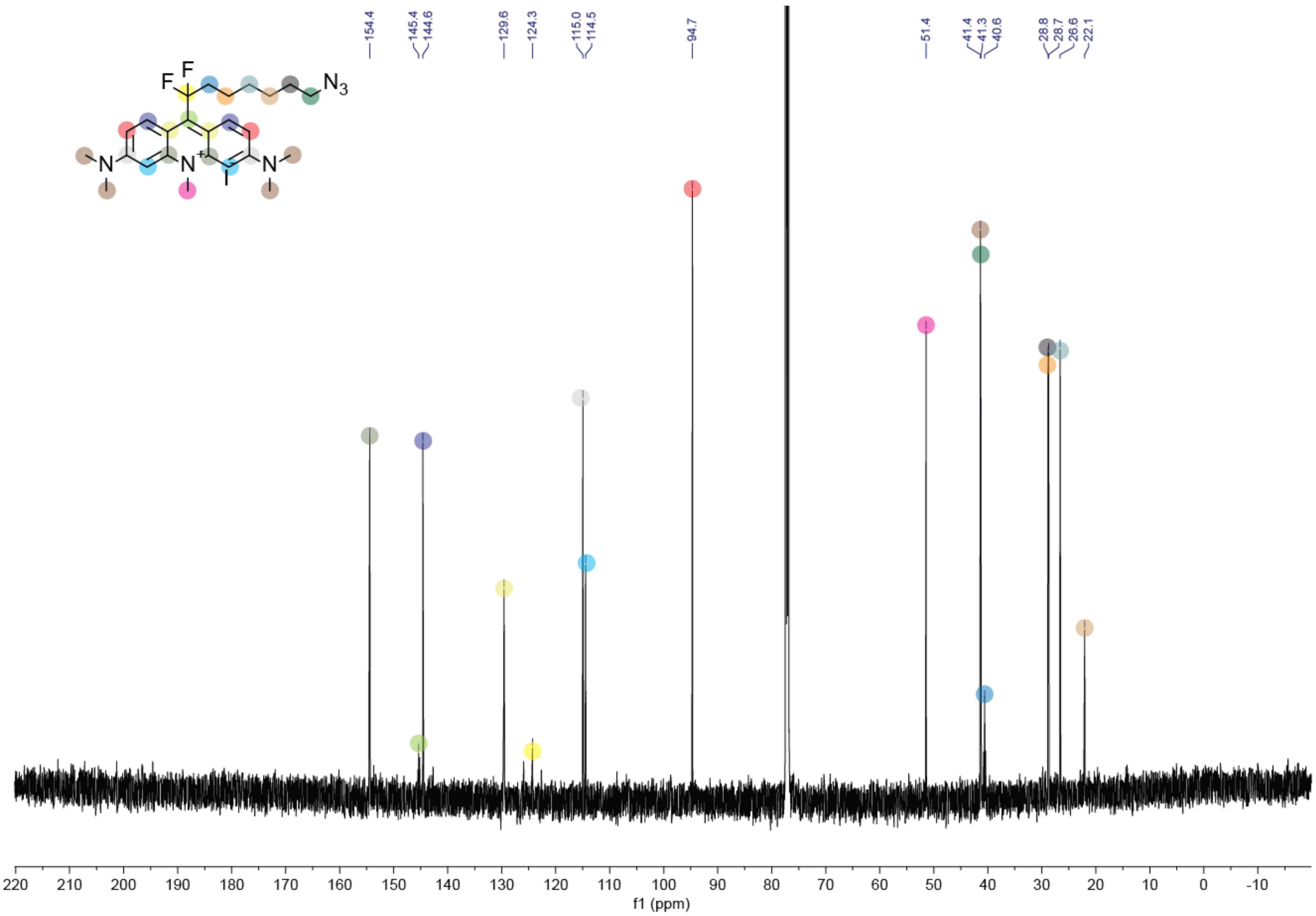

154.4
145.4
144.6
129.6
124.3
115.0
114.5
94.7
51.4
41.4
41.3
40.6
28.8
28.7
26.6
22.1
F
F
N3
N
N+
N
220 210 200 190 180 170 160 150 140 130 120 110 100 90 80 70 60 50 40 30 20 10 0 -10
f1 (ppm)

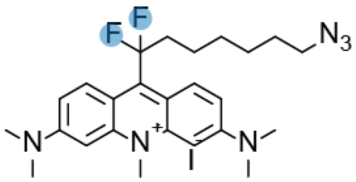
F
F
N3
N
N+
I
N

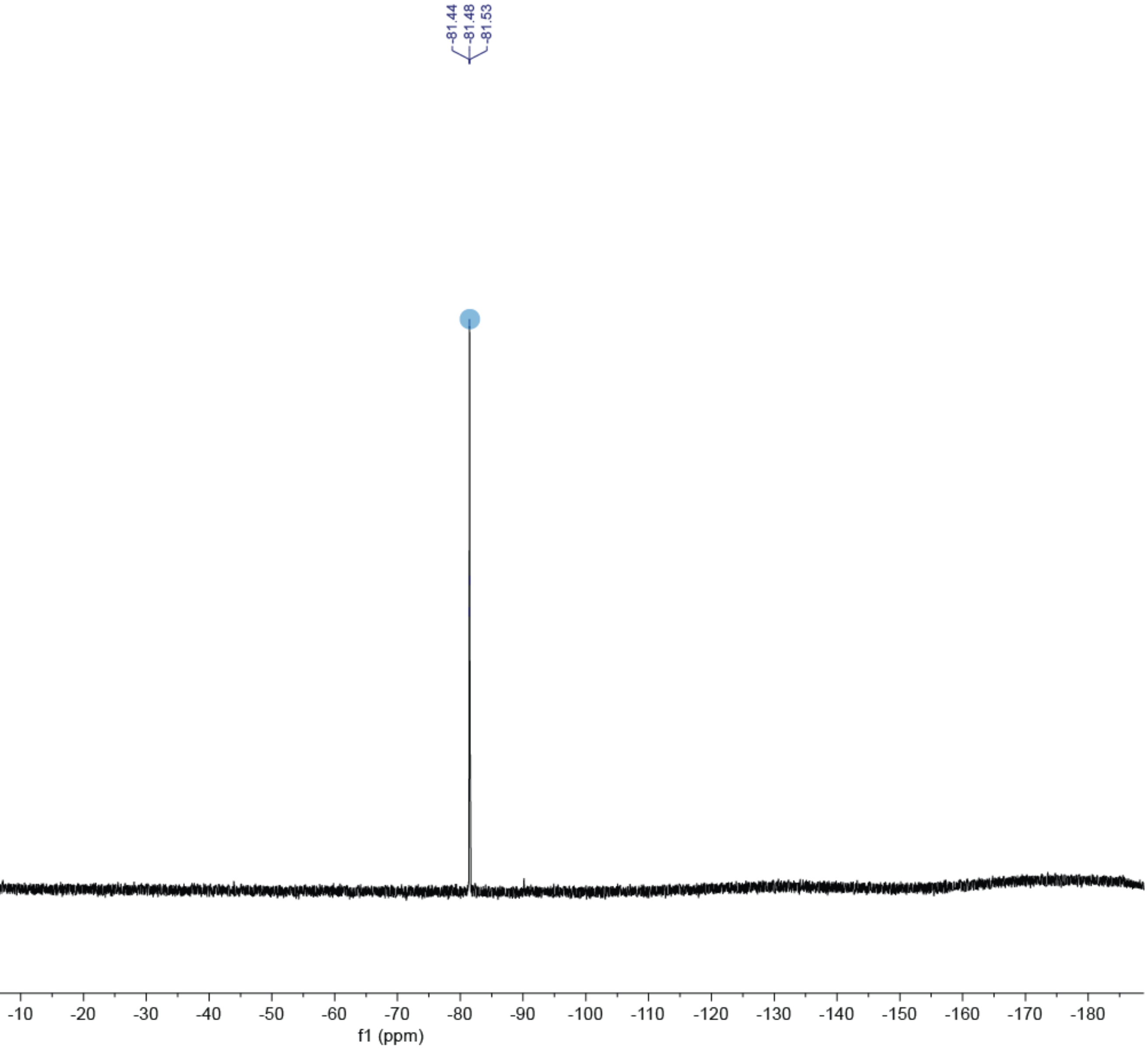
-81.44
-81.48
-81.53
50 40 30 20 10 0 -10 -20 -30 -40 -50 -60 -70 -80 -90 -100 -110 -120 -130 -140 -150 -160 -170 -180
f1 (ppm)

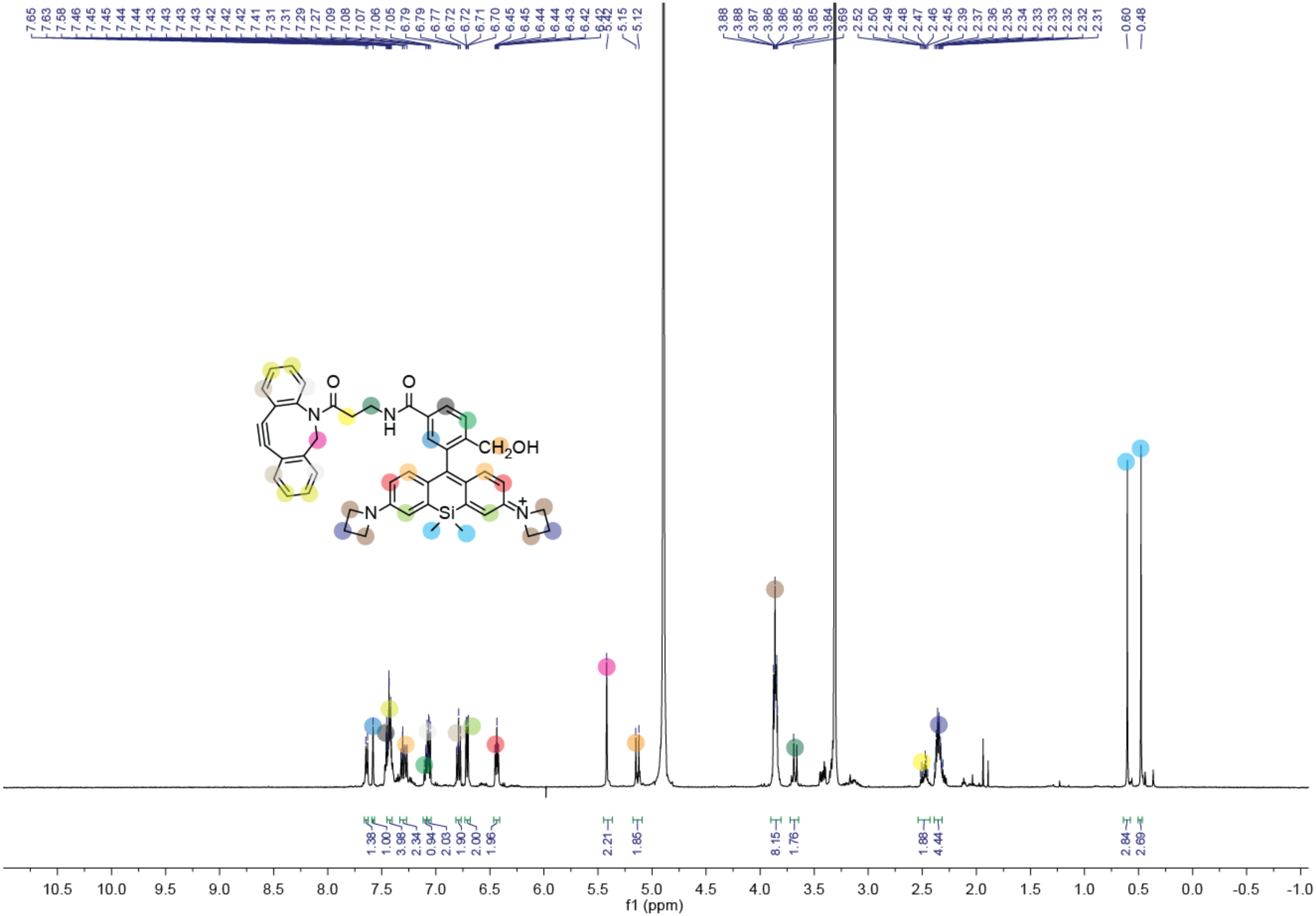

7.65 7.63 7.58 7.46 7.45 7.45 7.44 7.43 7.43 7.43 7.42 7.42 7.42 7.41 7.31 7.29 7.27 7.09 7.08 7.07 7.06 7.05 6.79 6.79 6.77 6.72 6.71 6.70 6.45 6.45 6.44 6.44 6.43 6.42 5.42 5.42 5.15 5.12
3.88 3.88 3.87 3.86 3.86 3.85 3.85 3.84 3.69 2.52 2.50 2.49 2.48 2.47 2.46 2.45 2.39 2.37 2.36 2.35 2.34 2.33 2.33 2.32 2.32 2.31 0.60 0.48
1.38 1.00 3.98 2.34 0.94 2.03 1.90 2.00 1.96 2.21 1.85 8.15 1.76 1.88 4.44 2.84 2.69
10.5 10.0 9.5 9.0 8.5 8.0 7.5 7.0 6.5 6.0 5.5 5.0 4.5 4.0 3.5 3.0 2.5 2.0 1.5 1.0 0.5 0.0 -0.5 -1.0
f1 (ppm)
O
O
N
N
H
CH2OH
N
Si
N+

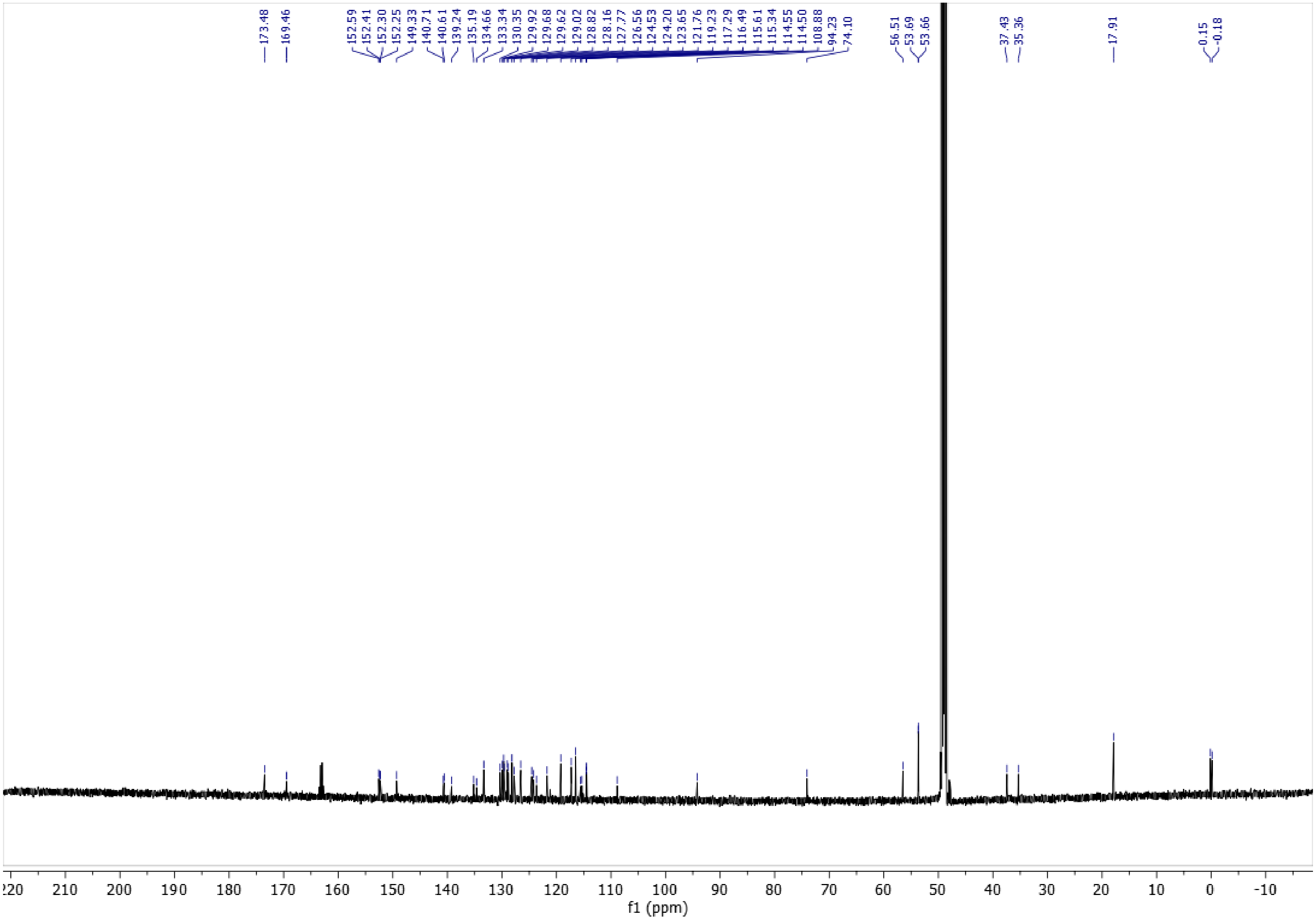

173.48
169.46
152.59
152.41
152.30
152.25
149.33
140.71
140.61
139.24
135.19
134.66
133.34
130.35
129.92
129.68
129.62
129.02
128.82
128.16
127.77
126.56
124.53
124.20
123.65
121.76
119.23
117.29
116.49
115.61
115.34
114.55
114.50
108.88
94.23
74.10
56.51
53.69
53.66
37.43
35.36
17.91
0.15
-0.18
220 210 200 190 180 170 160 150 140 130 120 110 100 90 80 70 60 50 40 30 20 10 0 -10
f1 (ppm)
